# The *Legionella pneumophila* peptidoglycan recycling kinase, AmgK, is essential for survival and replication inside host alveolar macrophages

**DOI:** 10.1101/2025.03.21.644609

**Authors:** Sushanta Ratna, Lina Pradhan, Marina P. Vasconcelos, Aastha Acharya, Bella Carnahan, Alex Wang, Arit Ghosh, Abigail Bolt, Jacob Ellis, Stephen N. Hyland, Ashlyn S. Hillman, Joseph M. Fox, April Kloxin, M. Ramona Neunuebel, Catherine Leimkuhler Grimes

## Abstract

Bacterial cells are surrounded by a dynamic cell wall which in part is made up of a mesh-like peptidoglycan (PG) layer that provides the cell with structural integrity and resilience. In Gram-positive bacteria, this layer is thick and robust, whereas in Gram-negative bacteria, it is thinner and flexible as the cell is supported by an additional outer membrane. PG undergoes continuous turnover, with degradation products being recycled to maintain cell wall homeostasis. Some Gram-negative species can bypass *de novo* PG biosynthesis, relying instead on PG recycling to sustain growth and division. *Legionella pneumophila* (hereafter *Legionella*), the causative agent of Legionnaires’ disease, encodes such recycling machinery within its genome. This study investigates the biochemical, genetic, and pathogenic roles of PG recycling in *Legionella*. Previously, we have shown that PG can be visualized in both model and native systems using a combination of *N*-acetylmuramic acid (NAM) probes and PG recycling programs. Here, two PG recycling gene homologs in the *Legionella* genome *lpg0296* (*amgK*) and *lpg0295* (*murU)* were identified and characterized; chemical biology strategies were used to rigorously track the incorporation of “click”-PG-probes. Deletion of *amgK* abolished PG labeling, while genetic complementation restored labeling. Additionally, copper-free click chemistry with ultra-fast tetrazine-NAM probes enabled live-cell PG labeling. The data suggest that *amgK* contributes to the pathogenicity of the organism, as *amgK* deletion increased *Legionella*’s susceptibility to antibiotics and significantly reduced *Legionella’*s ability to replicate in host alveolar macrophages. An intracellular replication assay demonstrated that while PG recycling is not essential for internalization, successful replication of *Legionella* within MH-S murine alveolar macrophages requires functional *amgK*. These findings underscore the essential role of AmgK in *Legionella*’s intracellular survival, emphasizing the importance of PG recycling in pathogenicity, and establish a foundation for developing novel *Legionella*-specific antibiotic strategies.

**Author summary:** In this work, we evaluated the functionality of the peptidoglycan kinase, AmgK, from *Legionella*. AmgK homologs from other organisms have been shown to play a role in peptidoglycan (PG) recycling. Here we performed a series of chemical biology, genetic and biochemical experiments to show that *Legionella* carries a functional AmgK. The data show that without AmgK *Legionella* becomes hypersensitive to Fosfomycin, an antibiotic that targets bacterial cell wall biosynthesis. Moreover, PG recycling is required for *Legionella* to successfully replicate within a host macrophage. Collectively, the data point to a role of PG recycling in pathogenicity and suggest that inhibiting AmgK could be a powerful strategy in treating Legionnaires’ disease.

## Introduction

Bacterial cells are surrounded by a dynamic cell wall that is essential for maintaining structural integrity. Peptidoglycan, a major component of the cell wall, consists of a repeating glycan backbone composed of *N*-acetylglucosamine (NAG) and *N*-acetylmuramic acid (NAM) linked by short peptide chains. This mesh-like network provides mechanical strength, enabling bacterial cells to withstand internal turgor pressure while supporting growth and division. In Gram-positive bacteria, the peptidoglycan layer is thick and highly cross-linked, while in Gram-negative bacteria, it is thinner, less cross-linked, and more flexible, positioned between the inner and outer membranes (1).

Despite subtle yet important differences in structure and regulation(1–3), the core biosynthetic steps are nearly identical across bacteria, including precursor synthesis in the cytoplasm, membrane-associated cytoplasmic-facing assembly, and membrane-associated periplasmic polymerization (1, 2, 4, 5) (Fig 1A).

**Fig 1.**
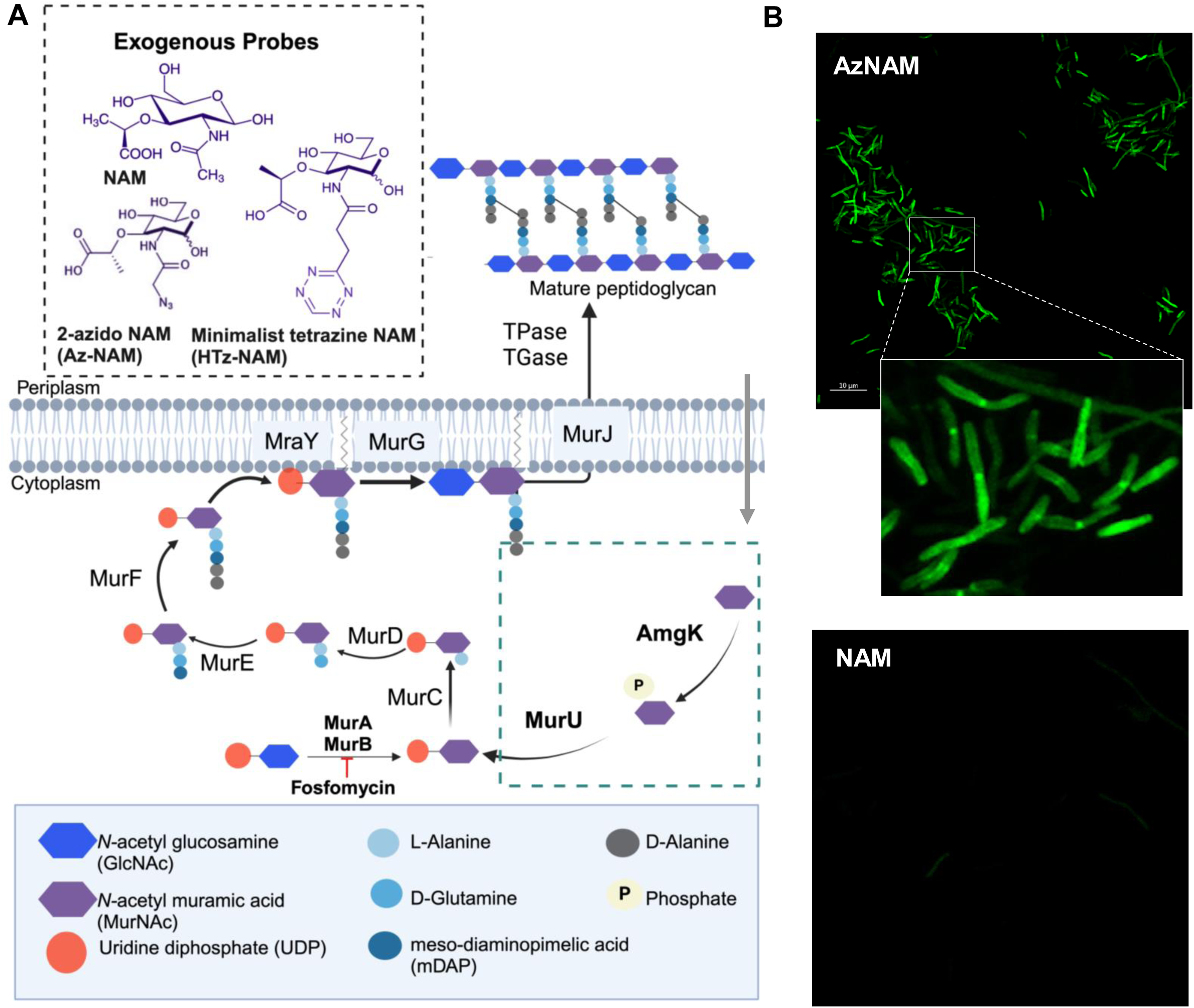
*Legionella* PG recycling pathway enables metabolic labeling of PG using bioorthogonal NAM probes. (A) *Structures* of NAM and NAM metabolic probes (Az-NAM) and (HTz-NAM) used to label *Legionella* PG*; Schematic* of peptidoglycan biosynthesis and recycling in *Legionella*. PG biosynthesis commences with MurA and MurB converting UDP-NAG in the cytoplasm to UDP-NAM. Recycling enzymes AmgK and MurU provide an alternative pathway, converting (NAM) into UDP-NAM. MurC-F extends UDP-NAM to Park’s nucleotide, which MraY transforms into Lipid I and MurG glycosylates forming Lipid II. MurJ transports Lipid II to the periplasm, where TGases and TPases cross-link the monomeric unit into mature peptidoglycan. Fosfomycin inhibits MurA, while exogenous probes such as Az-NAM and HTz-NAM enable the tracking of peptidoglycan synthesis. (B) *Representative* confocal micrographs showing *Legionella* cells grown with 6 mM AzNAM (top) or 6 mM NAM (bottom) for 3 hr, fixed, and conjugated with Alk-488. Data are representative of experiments performed in technical triplicate and biological triplicate.

The process of building PG begins with the enzymes, MurA and MurB, converting UDP-NAG to UDP-NAM, followed by the sequential addition of a pentapeptide chain by ligases MurC-MurF. MraY tethers the precursor to the membrane, MurG glycosylates this precursor with an NAG residue (forming Lipid II) and MurJ translocates the monomeric unit across the membrane (Fig 1A). The final PG structure is assembled as transglycosylases polymerize Lipid II, and transpeptidases cross-link adjacent peptide chains, ensuring cell wall integrity (6). Different bacterial species use a variety of linkage strategies in the peptide stem (3-3 vs 3-4), unusual amino acid side chains in the pentapeptide, including diaminopimelic acid (DAP), ornithine and iso-glutamine (iE). For iE, it should be noted that the pentapeptide chain is grown through the side chain and not the amino acid backbone. In addition, the percentage of cross-linking between different bacterial species can vary (7). Elegant studies in the early 1980s demonstrated that *Legionella*’s cell wall is highly crosslinked (8), permitting the cell to survive in high concentrations of lysozyme, an enzyme which cleaves the glycosidic linkages between NAG-NAM. Thus, these bacterial jackets have evolved to protect the organisms from the environments in which they live (9–11).

It should be noted that PG is not a static polymer. PG must undergo continuous remodeling during various cellular processes including growth, division, germination, and sporulation. Bacteria have developed mechanisms to recover and reuse its components rather than relying solely on *de novo* synthesis. PG biosynthesis uses ATP and NADH (Fig 1A); the process is anabolic and requires a significant amount of resources from the cell (12). Using *E.coli* model systems, PG recycling programs have been revealed (12, 13) and the community is beginning to learn the lengths that bacteria will go to recover these precious building blocks (14). Elegant work from the Mayer’s lab has shown, that *Pseudomonas spp*. can salvage NAM from degraded NAM-peptides using the recycling enzymes AmgK (*N*-acetylmuramate/N-acetylglucosamine kinase) and MurU (*N*-acetylmuramate α-1-phosphate uridylyltransferase) (13) (Fig 1A). This recycling pathway allows bacteria to bypass the conventional UDP-NAM synthesis via MurA and MurB (Fig. 1A), providing an alternative route for PG precursor generation and a resistance mechanism to the antibiotic, Fosfoymcin (Fig 1A). By recycling PG, bacteria can conserve energy and resources, supporting survival and adaptation in nutrient-limited environments.

Importantly, released PG fragments are well known to cause an immune response to the bacteria’s host (15). These small immunostimulatory fragments, generated by a variety of bacterial PG remodeling enzymes and host enzymes including lysozymes and muramidases (12) are recognized by innate immune receptors and part of the hosts innate immune defense systems (15–18). Thus, by restricting the release of immunostimulatory PG components, recycling provides bacterial pathogens with an added advantage in immune evasion.

The Gram-negative *Legionella pneumophila* (hereafter *Legionella*) is a facultative intracellular pathogen found in both fresh and potable water systems, where it parasitizes free-living amoebae. Upon inhalation of contaminated water droplets, *Legionella* replicates in alveolar macrophages, causing a severe pneumonia known as Legionnaires’ disease (19, 20). Following phagocytosis-mediated uptake, the bacteria remain enclosed in a replication-permissive membrane-bound compartment, known as the *Legionella*-containing vacuole (LCV), which avoids degradation by preventing fusion of the LCV with lysosomes (21, 22). To support its intracellular lifestyle, *Legionella* exploits host cellular processes by delivering bacterial effector proteins via the Dot/Icm Type IVb secretion system (23, 24). This specialized protein complex, spanning the inner and outer membranes, facilitates translocation of over 300 bacterial proteins, known as effector proteins, directly into the host cell. Dot/Icm-deficient bacteria fail to modify the phagosome for replication and are subsequently trafficked to the lysosome. The proper positioning of the Dot/Icm system at the bacterial poles is likely influenced by peptidoglycan dynamics. Consistent with this idea, studies have revealed that peptidoglycan editing, namely deacetylation of the glycan backbone, plays a crucial role in correctly positioning the Dot/Icm machinery at the bacterial pole (25, 26). Thus, localized PG turnover at the bacterial poles may facilitate the structural remodeling necessary for proper assembly and stabilization of the secretion machinery. However, the extent to which *Legionella* can recycle PG components has remained understudied. Homologs of key recycling proteins such as MupG, MurQ, and NamZ appear to be absent in *Legionella*, raising questions about its ability to recycle NAM (27). Interestingly, *Legionella* species do possess AmgK and MurU, leading us to hypothesize that these enzymes could allow *Legionella* to bypass *de novo* PG biosynthesis and facilitate PG recycling. We propose that this capability would enable *Legionella* to thrive in nutrient-limited environments, where NAG, NAM, and other essential PG biosynthesis cofactors are scarce.

In this study, we provide strong evidence that *Legionella* has the ability to recycle PG components. We investigated the role of the two key enzymes, AmgK and MurU, in *Legionella* PG recycling using chemical biology, genetic analysis, biochemical characterization, and infection assays. To assess PG biosynthesis, we employed NAM chemical biology probes with distinct biorthogonal features, permitting the tagging of specific cell wall features in complex cellular environments. Our lab has previously developed rigorous workflows to incorporate NAM probes directly into the PG (Fig 1A) (28). The ability to incorporate a chemical probe biochemically has been shown in a variety of bacterial settings, including *E. coli*, *B. subtilis*, *P. aeruginosa* and *S. aureus*, (28–31). In the work presented here, the metabolic uptake of the bioorthogonal PG probe 2-azido-N-acetyl-muramic acid (AzNAM) by *Legionella* cells was first verified using mass spectrometry and flow cytometry. Our findings demonstrate that AmgK and MurU are sufficient to drive PG recycling in *Legionella*, and deletion of *amgK* significantly reduces labeling and recycling efficiency. Although *amgK* and *murU* were not required for growth on charcoal-yeast-extract (CYE) medium, susceptibility to Fosfomycin increased ten-times in the absence of AmgK, compared to wild-type. Enzyme kinetics revealed that AmgK from *Legionella* exhibits activity ten-times than that of *P. putida*. Further analysis showed that AmgK is essential for intracellular survival and replication, indicating the critical role of PG recycling in bacterial persistence within host cells. Ultimately, this work provides key insights into how AmgK and MurU contribute to bacterial persistence within the host. Our findings suggest that targeting this recycling pathway could lead to novel therapeutic strategies for treating Legionnaires’ disease.

## Results

### Metabolic labeling with bioorthogonal NAM probes reveals peptidoglycan recycling in Legionella pneumophila

Facultative intracellular bacteria, capable of surviving both inside and outside host cells, encounter distinct metabolic and environmental challenges (32, 33). We hypothesized that PG recycling would be advantageous for these bacteria, supplying essential building blocks for cell wall repair and remodeling, thereby supporting their intracellular persistence. This may be particularly important for intravacuolar bacteria, such as *Legionella*, which are confined within a membrane-bound vacuole where nutrient availability may be tightly controlled by the host. Phylogenetic analyses have revealed the presence of AmgK and MurU homologs in the *Legionella pneumophila* genome (13, 27). Based on sequence similarity, *lpg0296* and *lpg0295* were identified as homologs of *P. putida amgK* and *murU*, sharing 39% and 46% identity, respectively (S1 Fig). Consequently, we set out to determine whether *Legionella* indeed utilizes the AmgK-MurU recycling pathway to funnel intermediates into the PG biosynthetic pathway.

To this end, a bioorthogonal labeling approach using azide-functionalized NAM (AzNAM) (Fig 1A), previously developed by our lab (28), was employed. Incorporation of AzNAM into the PG of growing cells were visualized by confocal micrscopy after fixation and conjugation of a fluorescent alkyne probe by copper (I)-catalyzed azide-alkyne click chemistry (CuAAC) (34, 35) (Fig 1B). The antibiotic, Fosfomycin, was added alongside NAM or AzNAM during labeling experiments. This antibiotic inhibits MurA, blocking *de novo* peptidoglycan biosynthesis, thereby forcing the bacteria to rely on the PG recycling pathways to maintain cell wall integrity (Figs 1A and 2A). Cells grown with the AzNAM analog exhibited a strong fluorescence signal, while control cells cultured with unmodified NAM showed no detectable fluorescence.

**Fig 2.**
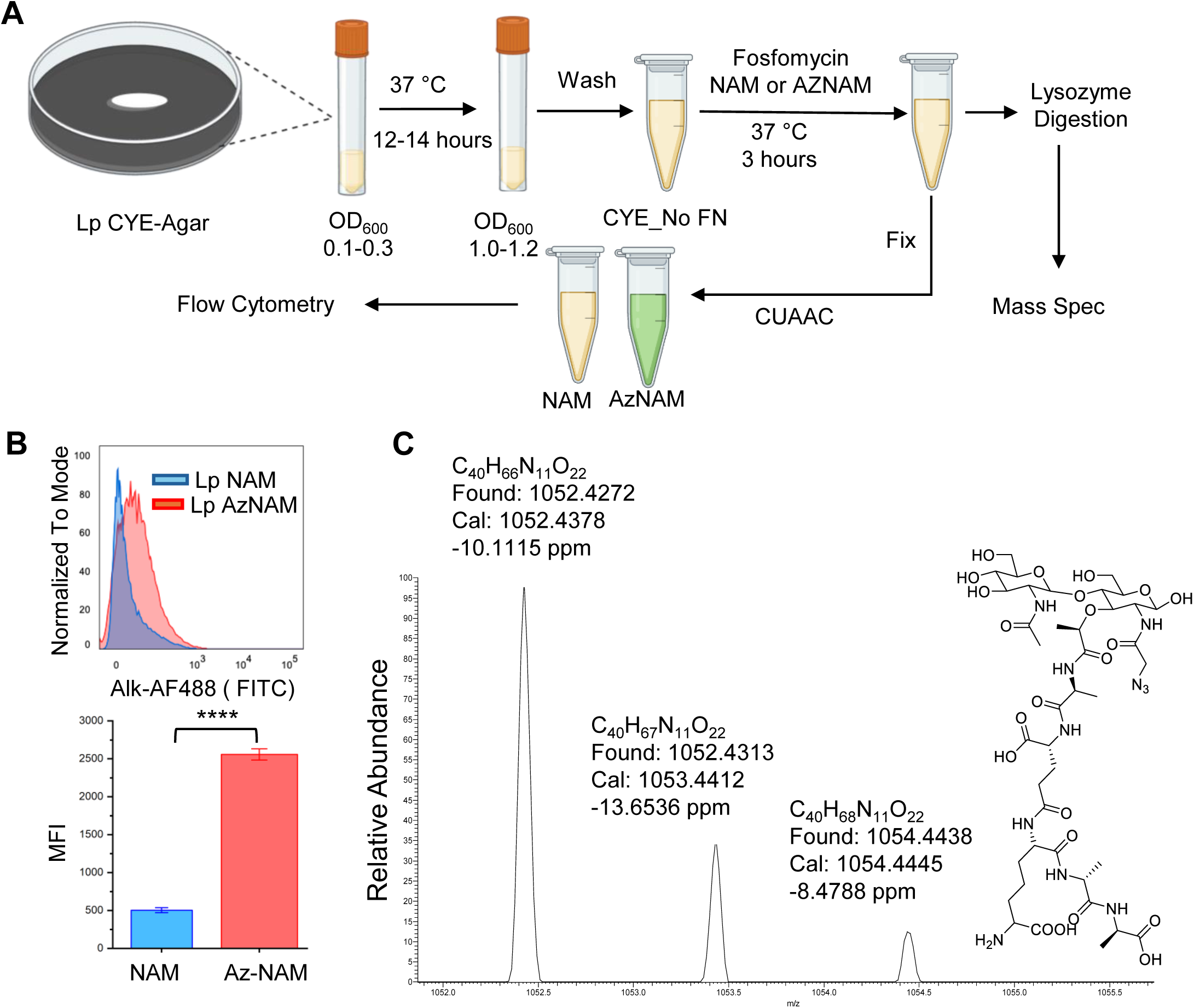
Validation of Peptidoglycan Labeling by Flow Cytometry and Mass Spectrometry. (A) Schematic workflow for both Flow Cytometry and Mass Spectrometry analysis of PG incorporation. (B) Flow cytometry analysis of cell wall labeling with Alk-488 in the presence of AzNAM or NAM. Histogram plot (left panel) and mean fluorescence intensity plot (right) show increased fluorescence signal intensity of cells labeled with AzNAM (red) compared to NAM control (blue). Statistical analysis was done by one way ANOVA with Tukey’s test using JMP pro 17 software (*\*\*\*\*p*< 0.0001). (C) Mass spectrometry analysis to verify AzNAM incorporation on lysozyme digested disaccharide fragments following remodeling with AzNAM. Data are representative of experiments performed in technical triplicate and biological triplicate.

To further quantify the extent of labeling, flow cytometry was performed, which revealed a five-fold increase in AzNAM-labeled cells relative to NAM-labeled cells. In addition, mass spectrometry data were used to confirm the incorporation of the exogenous AzNAM probe into the *Legionella* PG (Fig 2C). Briefly, bacterial cells were grown in liquid media until they reached exponential growth phase. At this time, cultures were supplemented with PG building blocks, either NAM or AzNAM, and incubated for an additional 3 hours. To confirm that the probe was directly incorporated into *Legionella*’s PG, remodeled cells were treated with lysozyme, which digests the PG into distinct NAG/NAM disaccharide subunits. The resulting fragments were analyzed via mass spectrometry. The data confirm the presence of a disaccharide containing the azide moiety, providing clear evidence that AzNAM probes were successfully integrated into the mature PG polymer, suggesting *Legionella* is capable of recycling PG using the NAM recycling pathway (Figs 1A and 2).

To determine the minimum effective concentration of AzNAM for labeling, varying concentrations of AzNAM were assayed, and a dose-dependent increase in fluorescence was observed; 0.75 mM AzNAM probe is sufficient for detectable incorporation into PG (S2 Fig). Notably, under these conditions, labeled cells retained their normal morphology and exhibited higher signal along the length of the bacterium as well as mid-cell at the division site where peptidoglycan synthesis is actively occurring during exponential growth (Fig 1B).

Collectively, these data (Figs 1 and 2) provide strong evidence that *Legionella* utilizes the AmgK and MurU recycling program (Fig 1A), during axenic growth. These finding demonstrate the utility of bioorthogonal NAM probes as a powerful tool for unraveling cell wall metabolism and remodeling in *Legionella*.

### AmgK is required for PG recycling in *Legionella*

*Legionella*’s ability to take up NAM and incorporate it into its PG, implicates AmgK as the kinase to facilitate this process (Fig 1). To test if AmgK is indeed responsible for channeling NAM into the PG biosynthesis, the dependency of PG labeling on AmgK was determined. An *amgK* deletion mutant was generated as well as a complemented strain carrying a plasmid encoded *amgK*. Two bioorthogonal labeling strategies were applied to determine NAM incorporation during growth of wild-type, *ΔamgK*, or the complemented *ΔamgK* strains. First, as described above, cells were cultured in media supplemented with AzNAM or NAM in the presence of Fosfomycin, fixed, and then labeled with Alk-488 to be visualized by fluorescence microscopy (Fig 3A). A robust fluorescence signal was detected for wild-type and complemented strains, but no fluorescence signal was observed for the *ΔamgK* mutant or in NAM-supplemented samples (Fig 3B).

**Fig 3.**
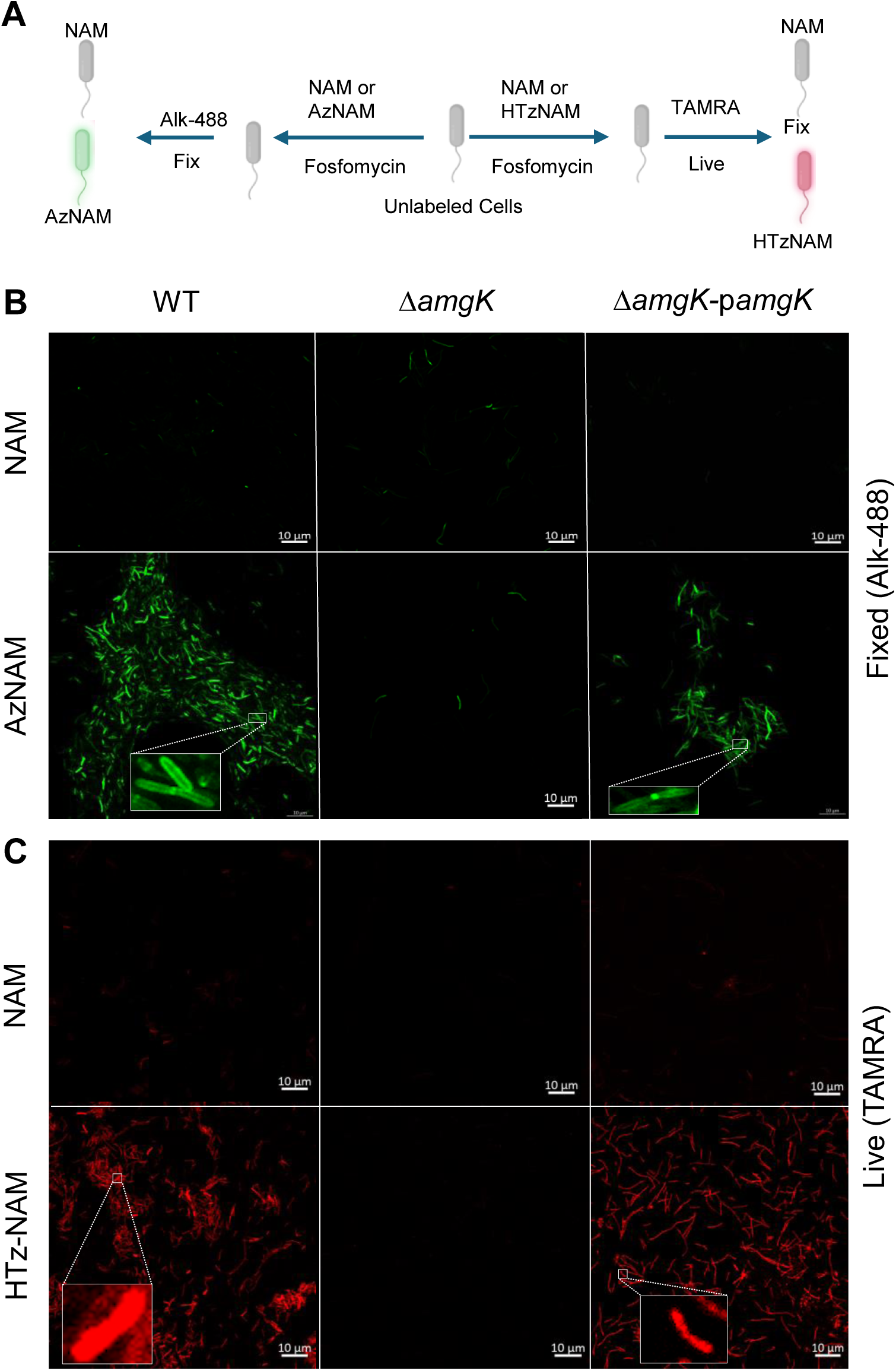
Metabolic labeling of *Legionella* requires the presence of AmgK. (A) Schematic representation of the two metabolic labeling techniques used to visualize peptidoglycan in *Legionella* strains. (B) Representative confocal micrographs of PG labeling in three *Legionella* strains: wild-type (Lp) (left), Δ*amgK* mutant (middle), and Δ*amgK* carrying the pMMB207c-*amgK* plasmid (right). For fixed-cell labeling, NAM was used as a negative control, while AzNAM served as the metabolic probe, reacting with Alk-488 to enable PG visualization. For live-cell labeling (C), NAM was again used as a negative control, while HTz-NAM was the metabolic probe, reacting with aTCO-TAMRA to label PG. *Each experiment was performed in at least three biological replicates*.

To further validate the role of AmgK in NAM incorporation, a second bioorthogonal labeling strategy using tetrazine ligation was employed (36). Tetrazine is a “click reaction”, which enables live cell labeling without the need for fixation (Fig 3C). For this approach, cells were cultured in media supplemented with HTz-NAM, a tetrazine-functionalized NAM analog (30), in the presence of Fosfomycin. Following incubation, cells were treated with aTCO-TAMRA, a transcyclooctene (TCO)-conjugated fluorescent probe, which reacts specifically with the tetrazine moiety via a rapid and highly selective tetrazine-TCO ligation reaction (37). The tetrazine-TCO ligation has been shown to have kinetics similar to enzymes and offers a key advantage over the traditional CuAAC click chemistry, which requires copper(I) catalysts that can be cytotoxic to bacterial cells (30, 38).

By using a copper-free approach, NAM incorporation was monitored in live *Legionella* cells, preserving their physiological state. Consistent with our previous findings using AzNAM (Fig 1B), a strong fluorescence signal was detected in wild-type and complemented strains, while no detectable labeling was observed in the *ΔamgK* mutant or in NAM-supplemented samples (Fig 3C), reinforcing the conclusion that AmgK is essential for funneling NAM back into the PG biosynthetic pathway.

The ability to monitor NAM incorporation in both fixed and live cells using distinct bioorthogonal labeling strategies underscores the robustness and versatility of the labeling approach. These results establish AmgK as a key enzyme responsible for incorporating NAM into *Legionella*’s PG. The data demonstrate that PG recycling in *Legionella* relies on the AmgK-dependent salvage pathway.

### Loss of AmgK increases susceptibility to Fosfomycin

As previously discussed, Fosfomycin (Fig 1A) inhibits the first committed step of peptidoglycan biosynthesis. To determine whether cell wall recycling contributes to Fosfomycin resistance in *Legionella*, the growth of wild-type was compared to a *ΔamgK* mutant strain using a viability assay. Cells were grown in liquid media for 6 hours and samples were then plated to enumerate the colony forming units (CFUs). In the absence of Fosfomycin, both wild-type and *ΔamgK* mutant strains exhibited comparable CFUs when plated on CYE media, indicating that the loss of *amgK* does not impact general growth under standard conditions (Fig 4).

**Fig 4.**
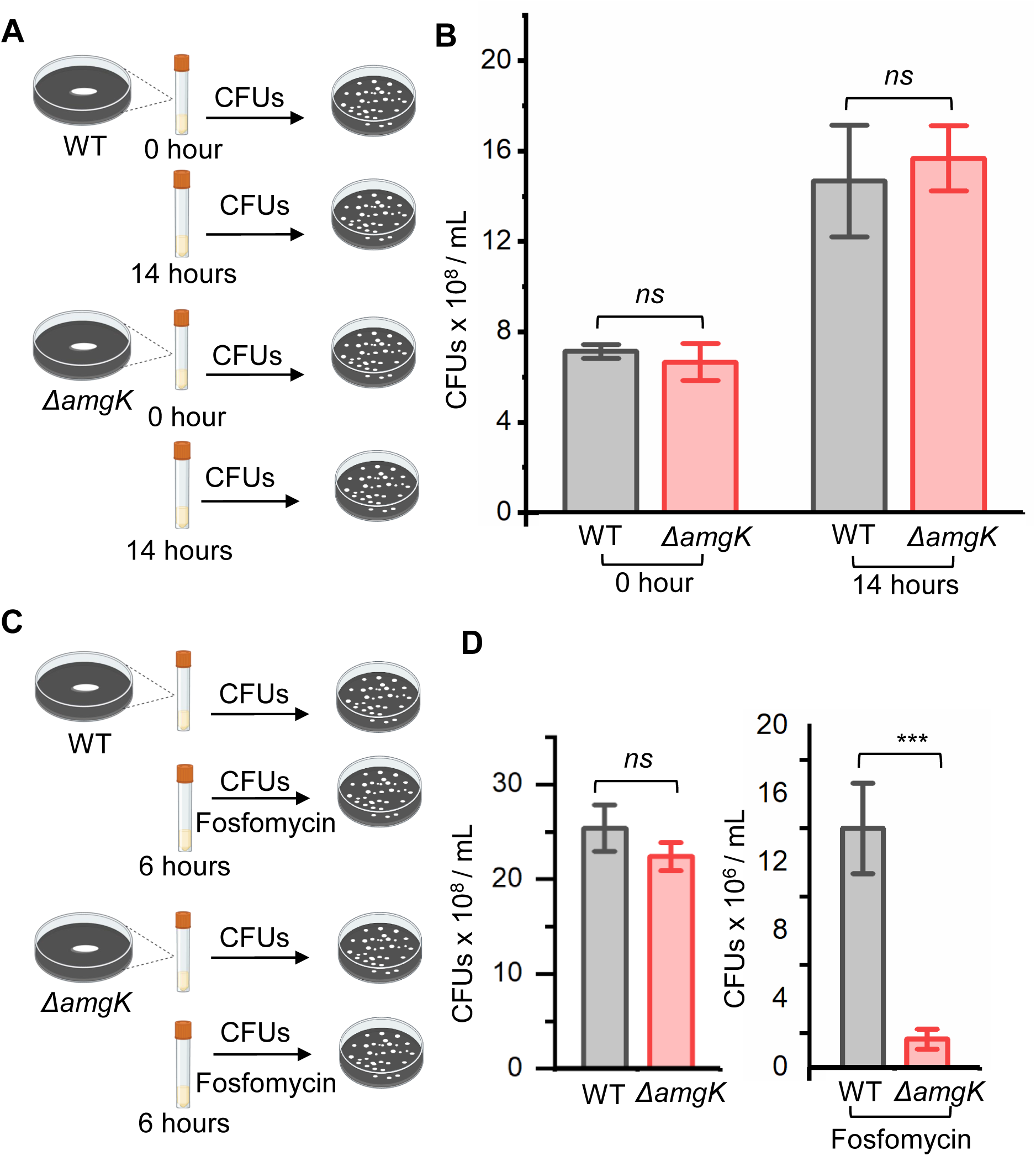
Effect of AmgK on cell growth and antibiotic resistance. (A) Schematic representation of the experimental approach for assessing cell growth. (B) Growth dynamics of *Legionella* in CYE medium with and without AmgK. (C) Methodology for evaluating the effect of Fosfomycin on *Legionella* growth. (D) CFU counts after 6 hours of incubation without or with Fosfomycin in CYE medium. All experiments were performed in three biological replicates; Error bars show +/- SD.

In contrast, when Fosfomycin was added to the media, both strains showed a significant reduction in CFUs compared to untreated cells, confirming the expected inhibitory effect of the antibiotic. However, the *ΔamgK* mutant displayed a markedly greater susceptibility, with a drastic decrease in CFUs after 6 hours of growth (Fig 4). The data indicate that AmgK plays a critical role in protecting *Legionella* from Fosfomycin-induced stress. These observations support the notion that PG recycling through AmgK is important for maintaining cell wall integrity when *de novo* PG biosynthesis is compromised.

### *Legionella* AmgK-MurU are unable to rescue growth in an *E. coli* NAM-dependent PG recycling model

We next sought to determine whether the *Legionella*-encoded AmgK and MurU are sufficient for NAM recycling. To assess this, we utilized an *E. coli* Δ*murQ* deletion mutant developed by Mayers and coworkers (13). Briefly, MurQ converts MurNAc-6P into GlcNAc-6P and is essential for the canonical PG recycling pathway in *E. coli*. Without MurQ, *E. coli* can no longer process MurNAc-6P into GlcNAc-6P, making it dependent on alternative pathways for PG recycling. This allows functional assessment of PG recycling enzymes from other bacteria, as has been shown previously (13, 28). To assess the functionality of the AmgK-MurU recycling pathway, the *amgK* and *murU* genes from *L. pneumophila* and *Pseudomonas aeruginosa* were introduced into *E. coli* Δ*murQ* using an expression vector; Two strains were generated: 1) *E. coli* Δ*murQ* expressing *Legionella* AmgK-MurU (EQ-LpKU) and 2) *E. coli* Δ*murQ* expressing *P. aeruginosa* AmgK-MurU (EQ-PaKU). Protein expression of AmgK and MurU in both EQ-PAaKU and EQ-LpKU lysates was confirmed by Western blot analysis (S6 Fig). Bacterial cultures were incubated for 2 hours and in the presence of absence of Fosfomycin. Cell density measurements show that without Fosfomycin, both strains exhibited normal growth, comparable to wild-type *E. coli* (Fig 5B). Conversely, in the presence of 200 µg/mL Fosfomycin, which blocks UDP-NAM synthesis, both EQ-LpKU and EQ-PaKU were unable to grow. However, growth was successfully restored when EQ-PaKU cells were supplemented with 6 mM NAM, demonstrating that UDP-NAM biosynthesis was rescued through the *P. aeruginosa* AmgK-MurU pathway. In contrast, growth of EQ-LpKU cells failed to recover despite NAM supplementation. This observation could be attributed to a number of factors including existence of unidentified components required for NAM salvage in *Legionella*, insufficient stability of the enzyme in *E. coli*, or lower catalytic efficiency of *Legionella* AmgK.

**Fig 5.**
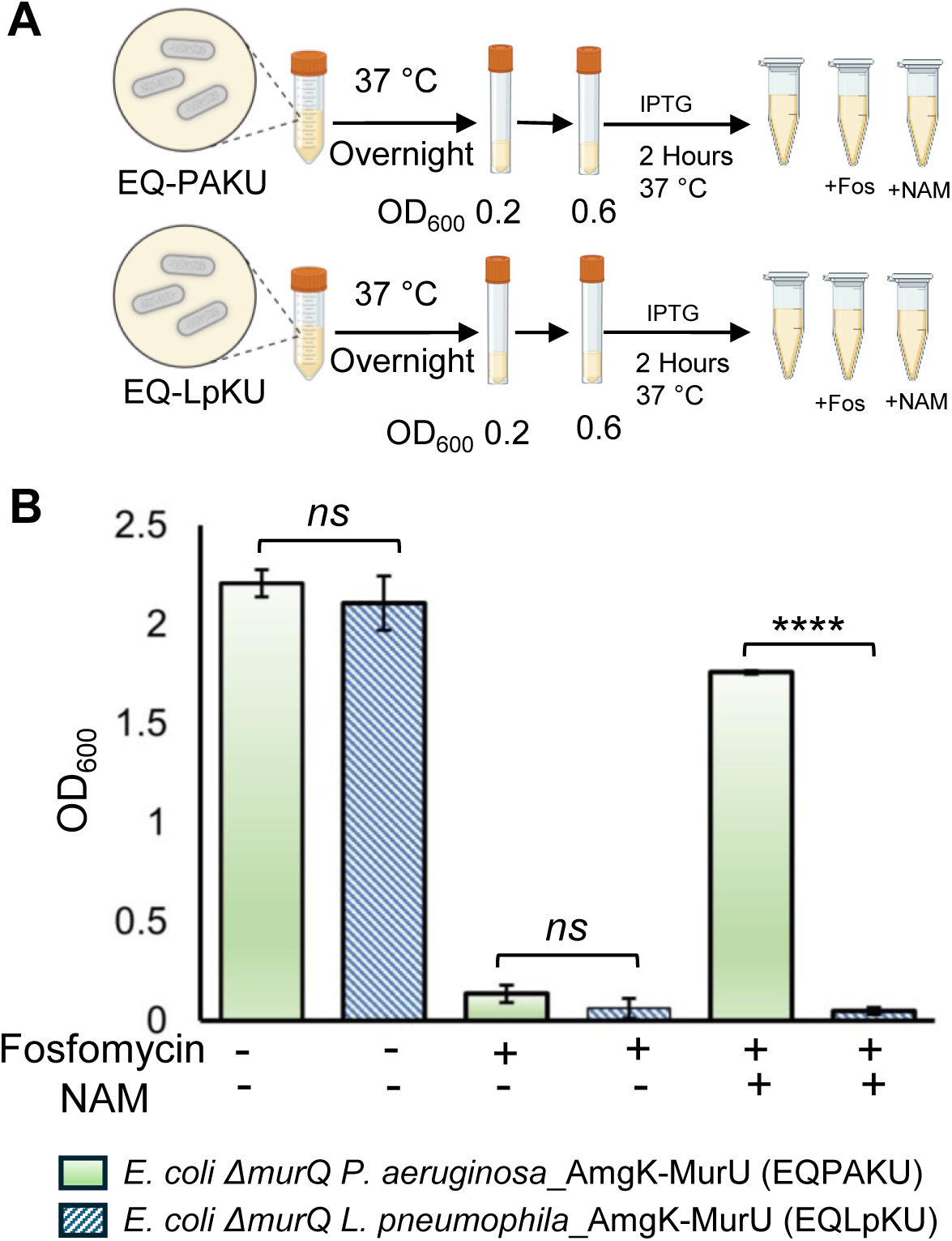
AmgK and MurU expression in *E. coli ΔmurQ* and survival assay. (A) Schematic representation of the viability assay. (B) Bar graph displaying cell density after two hours of growth under different conditions shown in the figure. Statistical analysis was performed on data from one experiment with three technical replicates, with the experiment conducted at least three times independently (\*\*\*\**p* < 0.0001).

### *Legionella* AmgK exhibits reduced catalytic efficiency

Both *E. coli* and *P. aeruginosa* have short doubling times (20–45 minutes), whereas *Legionella* grows significantly slower, with a doubling time of approximately 2–3 hours, even under optimal laboratory conditions (39, 40). Given this slower growth rate, it is reasonable to expect that *Legionella’s* AmgK may have adapted to function at a reduced catalytic pace, aligning with the organism’s extended doubling time. As a result, the dynamics of PG recycling in *Legionella* likely operate at a different tempo, with a reduced demand for rapid peptidoglycan turnover compared to fast-growing bacteria like *E. coli* and *P. aeruginosa*. To further explore this notion, we set out determine the catalytic efficiency of the *Legionella* AmgK.

First, we noted that the AlphaFold-predicted structures of AmgK from both species were highly similar and nearly superimposable (Fig 6A). To characterize the enzymatic activity, *Legionella* AmgK was expressed in *E. coli* as GST-AmgK and purified to 99% purity using a GST-resin column in the presence of PreScission Protease, yielding the expected 38 kDa band (Fig 6B). Kinetic analysis, using an ATP-coupled assay, revealed a K_m_ of (4.90 ± 0.0716) x 10^-5^ M, V_max_ = 4.757, *k*_cat_ = (2.370 ± 0.0717) s^−1^, and a catalytic efficiency of (4.84 ± 0.854) × 10^4^ M^−1^s^−1^ (Fig 6C). When compared to the previously published *P. putida* AmgK, the catalytic efficiency of *Legionella* AmgK is approximately tenfold lower (41). The diminished catalytic efficiency of *Legionella* AmgK lends support to the idea that *Legionella* may rely on a slower, yet sufficient, enzymatic turnover to meet its peptidoglycan recycling demands. This suggests that AmgK activity is tuned to the metabolic and replicative needs of bacterial species, reflecting broader evolutionary adaptations in PG recycling.

**Fig 6.**
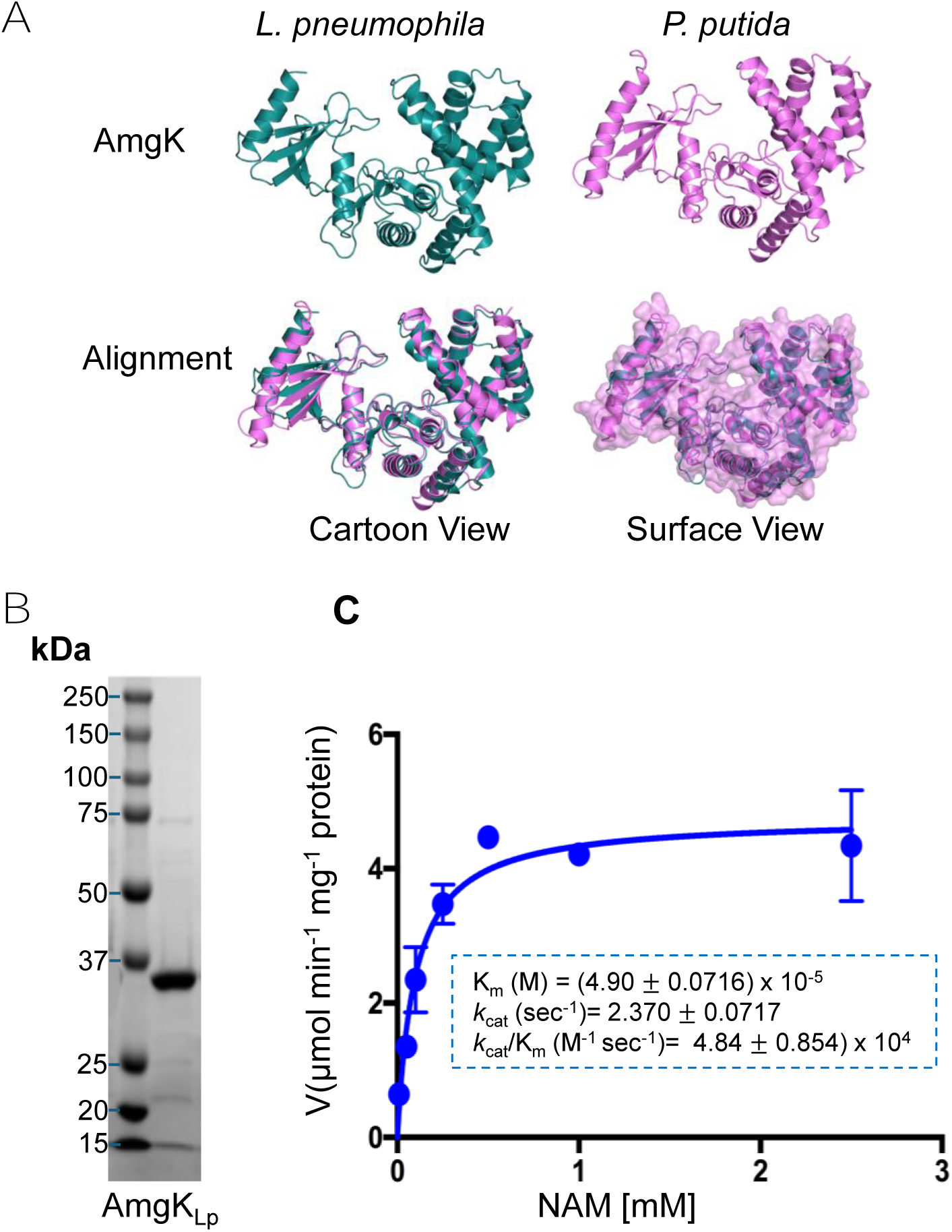
Structural and kinetic analyses show that *Legionella* AmgK is a peptidoglycan recycling kinase. A) Structural alignment of *Legionella* and *P. putida* AmgK, predicted using AlphaFold, shows high structural similarity between the two homologs. (B) Purification of *Legionella* AmgK using a GST-resin column, followed by cleavage with Prescission protease to remove the GST tag. SDS-PAGE confirms successful purification of untagged AmgK (∼38 kDa). (C) Kinetic parameters of *Legionella* AmgK determined using an ATP-coupled enzymatic assay, measuring substrate turnover and enzymatic efficiency.

### Bioorthogonally Labeled *Legionella* are Readily Phagocytosed by MH-S Macrophages

With the NAM labeling method rigorously validated in *Legionella*, we next sought to confirm that PG labeling preserves *Legionella* uptake efficiency into host cells. To this end, we assessed phagocytosis of *Legionella* by MH-S murine alveolar macrophages, a cell line regarded as a useful model for intracellular bacterial pathogens linked to respiratory diseases, including *Legionella pneumophila*, *Mycobacterium tuberculosis*, and *Chlamydia pneumoniae* (42–44). MH-S cells were challenged with AzNAM- or control unmodified NAM-labeled *Legionella*. At 30, 60, 90, and 120 minutes post-infection, cells were fixed, incubated with the Alk-488 probe in the presence of copper, and then visualized by confocal microscopy (Fig 7). Across all time points, *Legionella* remained robustly labeled and clearly visible within host macrophages. By 30 minutes, both labeled and control bacteria were internalized. Actin remodeling is a crucial step in *Legionella’s* infection, as the pathogen manipulates the host cytoskeleton to establish its replicative niche (45). By staining with phalloidin, which specifically binds F-actin, we observed that actin was indeed present at the LCVs at 2 hours post-infection (Fig 7). These findings support the notion that bioorthogonal PG labeling can be applied for tracking the early stages of *Legionella* infection within host cells and the probes developed in this study will be a useful resource for the field.

**Fig 7.**
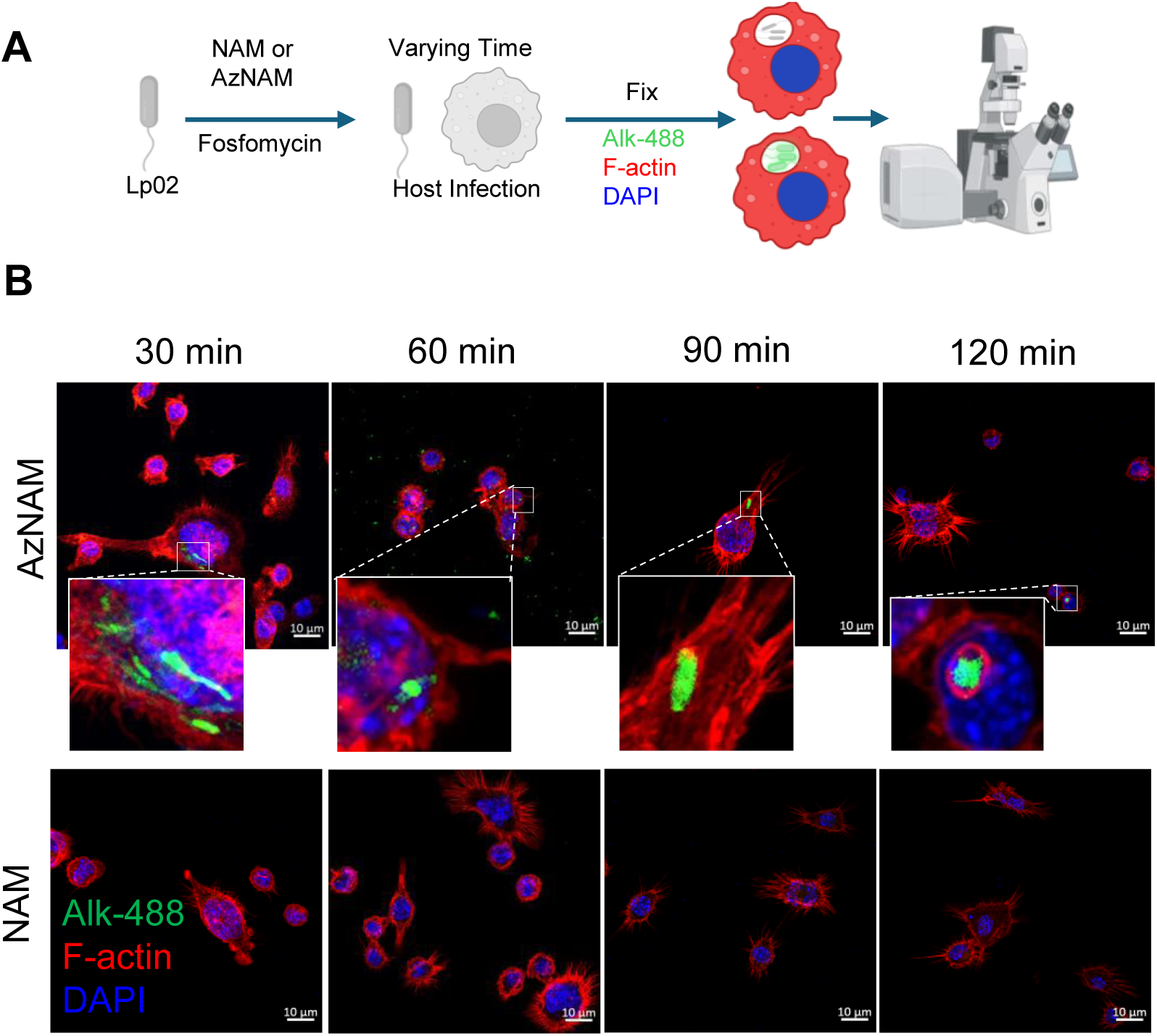
Infection study of MH-S invasion with *Legionella*. (A) Schematic depicting workflow for macrophage infection with labeled *Legionella*. (B) Representative confocal micrographs of MH-S cells infected with *Legionella* labeled with either AzNAM or NAM. Infected macrophages were fixed at 30, 60, 90, and 120 minutes post-infection, followed by CuAAC click labeling (*Legionella*, Alk-488, green), and subsequent staining with Phalloidin-TRITC (F-actin, red) and DAPI (nucleus, blue). Single-channel images of AzNAM-labeled *Legionella* inside MH-S and NAM-labeled *Legionella* inside MH-S are shown in Figs S8 and S9, respectively. The time course was conducted over at least three biological replicates; representative images from one time course are shown.

### AmgK is required for *Legionella* survival and replication within MH-S alveolar macrophages

To investigate the role of AmgK in *Legionella* intracellular survival, MH-S murine alveolar macrophages were challenged with a wild-type or a Δ*amgK* mutant. Both strains constitutively expressed *thyA* to circumvent thymidine limitation during intracellular replication. MH-S murine alveolar macrophages were challenged with the two strains at a multiplicity of infection (MOI) of 1, as previously described (46). At 2 hours post-infection, CFUs indicated that a similar number of intracellular bacteria was present in cells infected with either wild-type or Δ*amgK*. Thus, at this stage, wild-type and *ΔamgK Legionella* are comparable in their ability to infect macrophages, indicating that AmgK is not required for host cell entry or early survival (Fig 8B).

**Fig 8.**
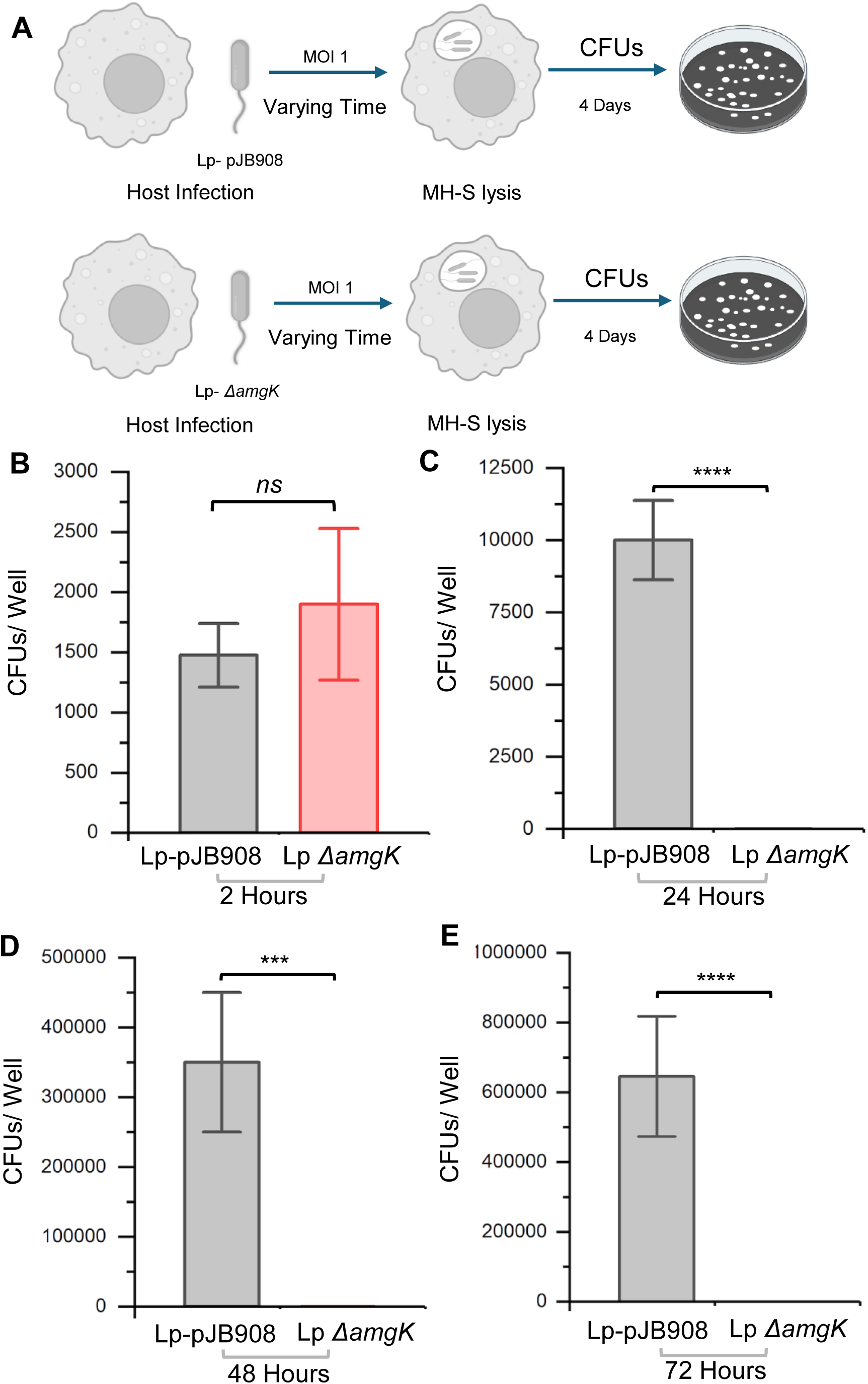
Impact of AmgK on *Legionella* Replication in MH-S Murine Alveolar Macrophages. (A) Schematic depicting workflow for *Legionella* infection and intracellular replication in MH-S alveolar macrophages, comparing strains with or without *amgK*. (B, C, D and E) Colony-forming unit (CFU) counts at four time points (2, 24, 48, and 72 hours post-infection), assessing bacterial survival and replication. Statistical analysis was performed on data from one experiment with three technical replicates, with the experiment conducted at least three times independently. (****p < 0.0001, *p < 0.001)

At all subsequent time-points sampled (24-72 hours), the wild-type continued to exhibit robust replication in macrophages, whereas the Δ*amgK* mutant showed no further growth and failed to persist, showing no detectable CFUs at later time points (Fig 8). Under normal circumstances, *Legionella* begins dividing within the LCV at 4-6 hours following infection. Thus, it appears that beyond the initial uptake into the LCV, *Legionella* requires AmgK to support intracellular growth. These findings suggest that while AmgK is not essential for initial infection, it is critical for *Legionella* intracellular survival and replication. The loss of AmgK likely disrupts PG recycling (Fig 1), which *Legionella* may rely on to maintain cell wall integrity under nutrient-limited conditions inside the host.

## Discussion

In Gram-negative bacteria, PG fragments released during cell wall remodeling can be salvaged through recycling. NAM recycling occurs via two distinct pathways: a catabolic route dependent on MurQ and an anabolic pathway mediated by AmgK and MurU (Fig 1). While some bacterial species encode both, others depend on a single route for PG recycling. First identified in *Pseudomonas putida*, the AmgK-MurU pathway bypasses the *de novo* synthesis of UDP-NAM, which is the first committed precursor of canonical PG biosynthesis (13, 47, 48). As a result, bacteria utilizing this pathway exhibit intrinsic resistance to Fosfomycin, an antibiotic that inhibits UDP-NAM biosynthesis (Fig 1A).

Here the identification and functional characterization of AmgK as a bonafide member of *Legionella’*s PG recycling machinery is reported through the use of chemical biology probes and genetics. These methods are then used to investigate the role of recycling enzymes in the pathogenicity of *Legionella*. The data clearly show that *Legionella* cells are able to incorporate a bioorthogonal probe such as AzNAM, into the PG. These data were rigorously validated using flow cytometry and mass spectroscopy analysis (Fig 2). This labeling strategy is important, as it is done in the context of the cell’s native environment. Previous work to metabolically/biochemically label PG on the muramic acid residues relied on genetic engineering to insert the recycling enzymes, *amgK* and *murU,* into the host genome such as *E. coli* (28, 30, 49). As *Legionella* already has the recycling enzymes, this step is unnecessary and opens the door to potentially use this methodology to detect *Legionella* in the environment.

Rigorous biochemical kinetic analysis showed that *Legionella’s* AmgK was capable of accepting a variety of probes, including alkyne and tetrazine derivatives (Fig 1) (29, 41). The alkyne and the tetrazine represent different types of “click” reactions. The azide is used with an alkyne counterpart in CuAAC, and in order to achieve fast rate constants, relies on a copper catalyst. Non-copper variations of CuACC exist, namely strain promoted alkyne azide click (SPAAC); this reaction takes advantage of ring strain in the alkyne system to negate the use of the copper catalyst, but reaction rates are orders of magnitude slower than CuACC. Tetrazines, which were used here, do not use copper to “click” to the transcycloocytne (TCO) counterpart and instead rely entirely on ring strain (30, 34–36, 50–61). Previously, tetrazine-TCO “click” reactions were not capable of labeling the PG using metabolic programs, as the tetrazine, traditionally a biphenyl tetrazine, was too large in size (41). However, recent developments of a minimal tetrazine NAM (HTzNAM, Fig 1A) enabled excellent labeling of PG (30). We were able to make use of this minimal tetrazine probe here to label *Legionella* PG. The flexibility in cell wall labeling of *Legionella* opens the door for future investigations of cell wall architecture and growth in this pathogen.

To complement the chemical biology approaches, we further investigated the function of *amgK* in the growth of *Legionella*. In order to study loss of function and gain of function of *amgK* gene, two strains were constructed: 1) *Legionella ΔamgK* and 2) complemented *amgK* gene in pMMB207c vector under lac promoter (S2 and S3 Figs). Importantly, both wild-type and complemented strains were capable of metabolic incorporation of NAM probes; whereas the deletion was not (Fig 3). These experiments confirmed the requirement of the *amgK* gene in PG recycling in *Legionella*, and indicate that this is the sole *amgK* homolog, as deletion of *amgK* obliterates labeling (Fig 3). We found that *Legionella*’s need for *amgK* changes in the context of the media used. In rich media, the deletion of *amgK* does not affect *Legionella* cells growth (Fig 4), as both wild-type and *ΔamgK* cells grow to similar CFUs .

Previous studies suggested that the presence of PG recycling machinery enzymes can affect the antibiotic resistance properties of cells (13, 48). Therefore, we investigated the effect of the presence or absence of *amgK* on the antibiotic Fosfomycin. Fosfomycin is a mimic of phosphoenolpyruvic acid, which is used by MurA in the PG biosynthetic pathway (Fig 1). MurA initiates the PG biosynthesis and as the data show, inhibition is deadly to the cell (Fig 4). AmgK enzymatic function is to phosphorylate NAM, negating the use of MurA. The results show that in the absence of Fosfomycin, both wild-type and *Legionella ΔamgK* cells grow normally. However, when Fosfomycin is present, only wild-type cells are capable of growing. These data suggest that *amgK* gives *Legionella* a fitness advantage.

AmgK was also investigated from a biochemical lens. Briefly, AmgK was overexpressed and purified in an *E. coli* system. With the purified enzyme in hand, kinetic assays were performed using the classic ATP coupled enzyme assay (41, 62). The data show that AmgK from *Legionella* exhibits kinetic properties suited to its growth needs (41). Given that *Legionella* is a slow-growing bacterium, particularly within the *Legionella*-containing vacuole (LCV), where nutrients are scarce (40), it is likely advantageous for the enzyme to operate at a more conservative pace, optimizing resource utilization in its intracellular environment.

In addition to the kinetic profile, the lack of other “traditional” recycling enzymes in *Legionella*’s genome warranted further investigation. To do this, we made use of a genetically encoded *E. coli* model system. When *amgK* and *murU* were initially characterized, Mayer and coworkers used a vector based strategy to introduce the genes into *E. coli ΔmurQ* (13). Their work showed that *amgK* and *murU* were sufficient to cause Fosfomycin resistance in a normally sensitive organism. To this end, *amgK* and *murU* from both *P. aeruginosa* and *Legionella,* were cloned into vectors and introduced into *E. coli ΔmurQ.* The data suggest that *P. aeruginosa amgK* and *murU* are sufficient to rescue when supplemented with NAM. Curiously, the same was not observed with *Legionella*, suggesting that in addition to kinetic limitations there may be other proteins working in concert with AmgK and MurU in the native environment to effectively recycle PG. The chemical probes presented here, computational analysis and genetics can be used in the future to unveil the complete PG recycling machinery in *Legionella*.

Finally, the importance of *amgK* in the pathogenesis of *Legionella* was investigated. It is well accepted that “nature does nothing uselessly” and here the question was raised regarding the importance of this PG recycling machinery in *Legionella*. Previous studies with known lung pathogens, including *M. tuberculosis,* showed different growth patterns and metabolic activity including exogenous fatty acid uptake and energy management if cells were grown in a similar condition as they would encounter inside alveolar macrophages (63, 64). These studies supported the hypothesis that *Legionella* might use a PG recycling machinery during intracellular replication as there would be a scarcity of available precursors to make *de novo* PG biosynthesis. Elegant work by Tamara and colleagues investigated the extent to which individual genes contribute to the fitness of *Legionella* during host infection (46, 65). Intriguingly, the absence of *amgK* decreased fitness of *Legionella* in the context of amoebae infection, but it seemed to increase fitness in the context of U937 macrophage infection (46). Remarkably, we found that *Legionella*’s ability to survive and replicate inside murine alveolar macrophages (MH-S cells) was dependent on *amgK.* Upon entry into the host cell, the internalized bacterium likely encounters nutrient-limiting conditions within the LCV, depriving it of essential building blocks for PG biosynthesis. Under these constraints, *de novo* PG synthesis may be downregulated to conserve resources, while PG recycling remains active, supporting critical processes such as PG remodeling. This study provides the first glimpse of PG biosynthesis in the LCV and highlights vulnerabilities that could be exploited for targeted drug development. Moreover, future work could reveal specific PG fragments that are important for stimulating the host-immune response (16, 66).

## Materials and Methods

### Cell Lines, Bacterial Strains, and Plasmids

The MH-S murine alveolar macrophage cells (ATCC CRL-2019^™^) were cultured using RPMI 1640 Medium (Corning) with 10% heat-inactivated FBS (Gibco) and 1% penicillin/streptomycin (Corning) at 37°C with 5% CO_2_ under sterile conditions.

A list of all bacterial strains, plasmids and cell lines used in this study is available in Table S1. *Legionella* strains in this study are derived from *L. pneumophila Philadelphia-1* Lp02 (*thyA*, *hsdR*, and *rpsL*). Lp02*ΔamgK* was generated by replacing the chromosomal *lpg0296* with *thyA*^+^ through allelic exchange as described below. *L. pneumophila* was grown in ACES (N-(2-acetamido)-2-aminoethanesulfonic acid) buffered yeast extract (AYE) liquid medium or on solid charcoal buffered yeast extract agar (CYE) medium. Both media were supplemented with 0.4 g/L iron(III) nitrate, 0.135 g/L cysteine, and 0.1 mg/mL thymidine for *thy−* auxotrophic strains.

*E. coli* strains were cultured in LB medium supplemented with 250 μg ml^−1^ chloramphenicol 250 µg/mL when appropriate. DH5α was grown in LB media. A summary of all plasmids and cells used in this work is available in Table S1.

### Construction of *Legionella ΔamgK* and *amgK* complementary strains

The pJB908 was used to PCR amplify the *thyA* gene and the pDG1662 (Bacillus Genetic Stock Center, BGSCID # ECE113) plasmid was used to amplify up and down stream nucleotide fragments for Allelic Exchange Substrate (AES). An in-frame deletion of *amgK* was obtained by allelic exchange removing all of the coding region except the sequences encoding residues 301-325. To generate the allelic exchange substrate (AES) DNA fragments were PCR-amplified with primers listed in S2 Table. To enhance AES transformation efficiency, DNA fragments from the pDG1662 vector were incorporated into the DNA construct, as described by Wu et al. (2019) (67). The pDG62 up (Adaptor_Up) and pDG62 down (Adaptor_Down) fragments were amplified with primers SR_F/pDG62_R and pDG62_F/SR_R, respectively, under conditions detailed in S3 Table. The *Legionella*_*amgK* upstream and downstream regions were amplified from genomic DNA, while the *thyA* gene was PCR-amplified from the pJB908 vector to allow for selection on CYE agar without thymidine. DNA fragments were gel-extracted (S3B Fig) and assembled using Gibson assembly as described in S3 Table. The assembled AES was then PCR-amplified with SR-F/SR-R primers (S3C Fig).

For transformation, *Legionella* cells were grown on CYE plates supplemented with thymidine (100 µg/mL), cysteine (400 µg/mL), and ferric nitrate (0.135 µg/mL) at 37°C for four days. A single colony was then streaked on a new plate and incubated at 30°C for 24 hours to form a patch. Then, 500 ng of AES in 10 µL was spread using a sterile glass rod and incubated for another 24 hours at 30 °C. Cells were harvested and streaked on a thymidine-free CYE plate and incubated at 37°C for four days. Colony PCR using Lp03-thyA Scr_Fwd and Lp03-thyA Scr_Rev primers was used to confirm the *amgK* gene knockout (S3D, S3E Figs). Replacement of *lpg0296* by *thyA* was verified via Sanger sequencing.

The pMMB207c-*amgK* complementation plasmid was generated by cloning *lpg0296* into pMMB207c-4xHA using Gibson assembly. Primers listed in S4 Table were used to amplify the *amgK* gene from genomic DNA. The pMMB207c-4xHA vector was linearized with *BamHI* and *HindIII* (S4A Fig). The gene and vector were assembled using HiFi 2x master mix under conditions specified in S5 Table and transformed into *E. coli* DH5α chemically competent cells. Transformants were selected on LB agar with chloramphenicol (250 µg/mL) (S4B Fig). Colony PCR confirmed the presence of *amgK*, followed by whole-plasmid sequencing for verification (S4C Fig). The *pMMB207c-amgK* plasmid was then introduced into *Legionella ΔamgK* as described earlier and selected on CYE plates with chloramphenicol (5 µg/mL) (S4D Fig). Colony PCR confirmed plasmid presence, which was further verified via whole-plasmid sequencing (S4E Fig).

Wizard Genomic DNA Purification Kit (Promega, Part # A1120) was used to isolate genomic DNA from *Legionella* and Qiagen plasmid miniprep kit for plasmid isolation from DH5α. PCR was done with Q5 high fidelity DNA polymerase (NEB, Cat #M0491S) and Phusion polymerase. PCR product was cleaned up using Qiagen kit or Zymeoclean Gel Recovery Kit (Zymo Research, Cat # D4001). Homologous genes were found using the Basic Local Alignment Search Tool (BLAST). NEBuilder® HiFi DNA Assembly Master Mix (NEB, Cat # E2621S) was used to do the Gibson Assembly. Reagents including Fosfomycin disodium salt (Sigma-Aldrich CAS # 26016-99-9), Isopropyl β-d-1-thiogalactopyranoside (IPTG) (GOLDBIO, CAS # 367-93-1), N-acetyl Muramic Acid (NAM) (Sigma-Aldrich, CAS # 10597-89-4), Phosphate buffered saline (PBS) (Corning, REF # 46-013-CM), Paraformaldehyde solution 4% in PBS (Santa Cruz Biotechnology, SC281692), CuSO_4_ (Fischer Scientific, CAS # 7758-99-8), BTTAA (7 mM stock) (Click Chemistry Tools), Sodium ascorbate (ACROS ORGANIC, CAS # 134-03-2), Fluorophore (Alk488, 1 mM stock) (Sigma-Aldrich), poly-l-lysine (Sigma-Aldrich, CAS # 25988-63-0), microscope slides (Fisher Scientific, Cat # 22037077) and ProLong Glass Antifade Mounting medium (Thermo Fisher Scientific, REF P36980) were used for bacterial cell wall labeling along with Az-NAM, which was synthesized in the Grimes lab. Tris (Hydroxymethyl) aminomethane (J&K, CAS # 77-86-1), Sodium Chloride (Thermo Fisher, CAS # 7647-14-5), EDTA (Sigma-Aldrich, CAS # 60-00-4) and Lysozyme (Sigma-Aldrich, CAS # 12650-88-3) were used to digest PG for mass spectrometry analysis.

### Metabolic labeling of *Legionella* with AzNAM probe

A fresh plate was prepared from 40% glycerol stock of *Legionella* in ACES-agar plate with supplemental nutrients by incubating at 37°C for four days to get a single colony. *Legionella* cells were grown in ACES broth (supplemented Thymidine (100 µg/mL), Cysteine (400 µg/mL) and Ferric Nitrate (0.135 µg/mL) with a serial dilution ranging OD_600_ 0.1 to 0.3 from a patch of a single colony that was grown for two days at 37°C in ACES-agar plate. Cells were incubated at 37°C with shaking for 12 h to 14 h to reach OD_600_ 0.9 to 1.4. The labeling protocol was adapted from our previous publications (28, 29, 31). In summary, 600 µL of overnight grown culture with OD_600_ 0.9 to 1.4 was collected by centrifugation at 8000 rpm (Eppendorf tabletop centrifuge, 6000g) for five minutes. Cells were washed with 600 µL ACES growth medium without Ferric Nitrate on it. We note that careful consideration of media components is essential, as the ferric nitrate supplement in AYE medium can reduce the AzNAM probe, diminishing labeling efficiency (S10 fig). Cells were resuspended in 190 µL ACES medium (without Ferric Nitrate). Then 6 mM NAM or AzNAM and 200 µg/mL Fosfomycin were added. Cells were incubated at 37 °C for 3 hours while shaking in the incubator. Cells were then collected and washed twice with 600 µL sterile 1x PBS at 10,000 rpm for two minutes and resuspended in 200 µL 4% paraformaldehyde in PBS for 20 minutes at room temperature to fix the cells. Cells were then resuspended on 190 µL PBS after washing twice at 300 µL PBS (10,000 rpm, 2 min). For click reaction sequentially added 1 mM CuSO_4_ solution, 140 µM BTTAA, 1.2 mM freshly prepared sodium-L-ascorbate in water and 0.8 µL of 1mg/mL stock of Alk-488. Then the mixture was incubated at room temperature for 30 minutes in a shaker at dark. Cells were washed 3 times with PBS and resuspended on 200 µL of PBS and incubated at room temperature for 60 minutes. Cells were washed once more and then resuspended on 100 µL PBS. 20 µL of cells were spread on a poly-l-lysine coated slide and incubated at room temperature for 30 minutes at dark. Then the cells were mounted with an antifade mounting medium and incubated overnight at room temperature before sealing it with clear nail polish. Cells were then visualized using Zeiss LSM800 microscope.

### Labeling efficiency based on probe concentration

To determine the optimal concentration of the NAM probe, cells were metabolically labeled with AzNAM across a concentration range (0.75 to 6 mM) (S2 Fig). The results demonstrate efficient labeling of *Legionella* PG, with 6 mM being most optimal for visualization of the PG using confocal microscopy. However, lower probe concentrations were still effective, indicating that the PG biosynthetic machinery readily incorporates and tolerates the AzNAM probe, allowing for efficient labeling of the cell wall (S2 Fig).

### Flow cytometry

Cells were labeled with AzNAM or NAM as described earlier (31). Cells were then diluted in PBS and flow cytometry was performed on BD FACS Aria Fusion High Speed Cell Sorter. Samples were briefly vortexed before each run. 50,000 cell counts were collected for each sample and were analyzed in triplicate, and fluorescence intensities (FITC-H, PMTV 434) for each sample were collected. Any potential autofluorescence from samples was collected and gated out via APC channel (PMTV 465). Histograms for control (*Legionella* NAM) vs *Legionella* AzNAM were overlaid (normalized to mode) on FlowJo^TM^.

### Mass spectrometry AzNAM probe incorporation assay

Probe incorporation assay was adapted from a previously published method (31). In short, 3 mL of *Legionella* cells were grown to OD_600_ 3.2-3.4 and washed with 1 mL of AYE medium supplemented only with thymidine (100 µg/mL), and cysteine (400 µg/mL). Cells were resuspended on 1 mL of the same medium supplementing with 50 µL of NAM/AzNAM (0.1 M stock) and 20 µL of Fosfomycin (10 mg/mL) and grew for 3 hours at 37°C. Cells were washed twice with PBS and then resuspended in 2.5 mL of digestion buffer (250 µL of 1 M Tris pH 7.9, 25 µL of 5 M NaCl, 20 µL of EDTA and 4.75 mL of sterile DI water). Freshly prepared 20 µL of lysozyme (50 mg/mL) were added in approximately a 12-hour interval for three days. Samples were then filtered with Amicon Ultra 3K filter device and lyophilized sample for overnight. Lyophilized samples were then resuspended on 5 times the volume of water by weight and submitted samples to orbitrap mass spec for analysis.

### Effect of AmgK protein on cell growth and antibiotic resistance

Wild-type and *ΔamgK* cells were grown by serial dilution as described earlier. For overnight preparation, initial dilution was maintained the same for both cells by taking OD_600_ 0.2-0.3 which is considered 0 hours. Then both cultures were incubated at 37°C for 14 hours. Both cells of 0 hours and 14 hours were diluted and plated to CYE-agar medium with appropriate supplements to get colony count. Fosfomycin concentration was determined based on the previous study (68). To check the effect of AmgK protein in term of Fosfomycin resistance, both wild-type and *ΔamgK* strains grown overnight were diluted in fresh AYE medium and incubated with 16 µg/ mL of Fosfomycin for 6 hours. After 6 hours of incubation, cells were diluted in sterile PBS and plated in CYE-agar medium to get the colony count.

### AmgK protein expression vector construction, protein expression and purification

AmgK was inserted into pGEX vector with GST tag at N-terminal under lac operon with HiFi assembly (NEB, NEBuilder Hifi DNA Assembly Cloning Kit, E2621S). Protein was then expressed and purified based on the method described previously (69) (S7 Fig). Primers were adapted from previous publications as mentioned in S8 table (62, 69, 70). pGEX-6p-1 vector was linearized with primers (S8 Table). *amgK* was amplified with appropriate overhang and HiFi assembly (S9 Table) was performed to insert gene under IPTG inducible lac operator with GST tag at the N-term of the protein with a precision protease cleavage site in between GST and respective genes (S7 Fig). Gibson assembled vectors and genes were transformed in chemically competent BL21 cells and selected in the presence of ampicillin/ carbenicillin (NEB, NEBuilder Hifi DNA Assembly Cloning Kit, E2621S). Whole plasmid sequencing was performed to verify the vector constructs.

A 10 mL overnight of BL21 cell culture was inoculated into 1 L of fresh LB medium supplemented with 100 ug/mL of carbenicillin antibiotic and incubated at 37°C until OD_600_ of 0.6 for both AmgK and MurU. The expression of GST-tagged protein was induced with 1 mM isopropyl-1-thio-β-D-galactoside (IPTG) at 16°C for 18-20 hours. Induced cells were harvested by centrifugation (8000 rpm, 15 minutes). A 1 L pellet protein was resuspended in 20 mL lysis buffer (50 mM Tris-HCl, pH 7.0, 150 mM NaCl, 2 mM DTT, 10% glycerol, 0.5% Triton X-100, and one complete mini EDTA-free protease inhibitor cocktail tablets from Sigma-Aldrich). Cells were sonicated (Total 2:59 min, 9.9 second pulse on, 39.9 second pulse off with 41% amplitude) and then centrifuged at 17000 rpm for 30 minutes. The supernatant was loaded onto a protein purification column with Glutathione Sepharose 4 B Fastflow beads (Cytiva) and incubated at 4°C for 3 h. The flow through was released and the column was washed twice with 10 mL of TBS Buffer (50 mM Tris-HCl pH = 7.0, 150 mM NaCl, 1 mM DTT, 1 mM EDTA, and 10% glycerol), 4 times with 10 ml GST wash buffer (50 mM Tris-HCl pH =7.0,(200, 300, 400 and 500 mM) NaCl, 1 mM DTT, 1 mM EDTA, and 10% glycerol), and then another 2 times wash of 10 mL of TBS buffer. 10 mL GST elution buffer (50 mM Tris-HCl pH = 7.0, 150 mM NaCl, 1 mM DTT, 1 mM EDTA, 10% glycerol, and 600 μL of Prescission protease) was added to the column and incubated for 16 hours at 4°C. AmgK elutions were collected with 10 mL of elution buffer and three subsequent elutions were performed with 5 mL elution buffer by shaking for 5 minutes each time. Protein samples were then concentrated using a 10,000 Da Amicon MWCO column. For long-term storage both protein samples were flash-frozen in glycerol with liquid nitrogen and stored at −80°C.

### Labeling Legionella, Legionella ΔamgK and Legionella ΔamgK-pMMB207c-amgK

Both *Legionella* and *Legionella ΔamgK* cells were labeled with AzNAM as described in the earlier section. *Legionella ΔamgK*-pMMB207c-*amgK* were grown in ACES-medium supplemented with ferric nitrate and cysteine without any antibiotic on it. Cells were grown with a serial dilution with the initial OD_600_ of 0.1 to 0.3 in ACES medium with 0.5 mM IPTG for 14 to 16 hours to reach OD_600_ of 0.9-1.3. Cells were washed with ACES medium and incubated with fresh ACES supplemented with 0.5 mM IPTG and cysteine without ferric nitrate on it. All the remaining steps are the same as previously described.

### Live labeling of *Legionella*, *Legionella ΔamgK* and *Legionella ΔamgK*-pMMB207c-*amgK* with HTz-NAM probe

Live labeling of *Legionella*-02 was performed according to the published protocol(30). Briefly, both wild-type and *ΔamgK* cells were grown with HTz-NAM as described in the earlier section. *Legionella ΔamgK*-pMMB207c-*amgK* were grown in ACES-medium supplemented with ferric nitrate and cysteine without any antibiotic on it. Cells were grown with a serial dilution with the initial OD_600_ of 0.1 to 0.3 in ACES medium with 0.5 mM IPTG for 14 to 16 hours to reach OD_600_ of 0.9-1.3. Cells were washed with ACES medium and incubated with fresh ACES supplemented with 0.5 mM IPTG and cysteine without ferric nitrate on it. All three cells were incubated for three hours with 6 mM of HTz-NAM. After three hours of incubation, 0.8 µl of aTCO-TAMRA (1mg/mL) were added to each tube and incubated for another 15 minutes at 37°C. Then washed and fixed cells with 4% PFA in PBS for 15 minutes. Then cells were washed 4 times and prepared slides with poly-L-Lysine coated glass. Images were taken with LSM 800 with 63x lens.

### Application of *Legionella* AmgK and MurU in PG recycling in *E. coli* Model system

The *amgK* and *murU* genes were cloned in pBBR MCS1 vector with HiFi assembly following protocol described in the earlier section. In short, FLAG tag was inserted to make recombinant *amgK* and *murU* for *Legionella* and HIS tag was added *P. aeruginosa* with primers and conditions described (S5 and S6 Tables). HIS tag inserted *amgK* and *murU* of *P. aeruginosa* were inserted in spectinomycin selectable antibiotic gene under IPTG inducible lac promoter (S6A Fig) while FLAG tag added *Legionella*’s *amgK* and *murU* were inserted in chloramphenicol (CamR) selectable antibiotic resistance gene (S6C Fig). Vectors were then inserted in the E. coli *ΔmurQ* model system. A subculture from overnight was diluted to OD_600_ 0.2 and incubated at 37°C until it reached OD_600_ 0.6. Then 1 mM IPTG was added and incubated for another 4 hours. Cells were then resuspended and lysed with 1.2 mL of B-PER bacterial protein extraction reagents (Thermo scientific, REF 78248). Cells were then sonicated using a bath sonicator for 3 minutes with one minute on and one minute off (on ice). Lysate was then centrifuged at 8000 rpm for 5 minutes and protein concentration was measured with Bradford reagent. 10-20 μg of protein was loaded in the SDS page and transferred into nitrocellulose paper. Then it was incubated by shaking with 10% skim milk in a transfer buffer (TBST) for one hour. The paper was washed 3 times with TBST buffer and resuspended in 1:1000 anti-FLAG/ 6x-HIS (Monoclonal ANTI-FLAG M2 (SIGMA) and HIS (H8 Invitrogen) both were produced on Mouse) in 1% skim milk in TBST buffer and incubated at 4°C on a rocker overnight. Then the paper was washed 3x with TBST buffer and added 1: 3000 IgG (produced in mouse) HRP in 1% skim milk in TBST and shaken at room temperature for one hour. Samples were washed 3x with TBST and then used Super Signal reagents for gel imaging. PA-KU and Lp-KU expression with HIS were verified with protein specific size in the gel (S6B and S6D Figs). Then live-dead assay was performed according to previously published methods (28, 29, 31). In short, overnight cultures were diluted to OD_600_ 0.2 and incubated until it reached OD_600_ 0.6. 1 mM IPTG was added when the cell reached OD_600_ 0.6. Cells were then grown for two hours by dividing three experimental categories where cells were grown in just LB medium, with 200 µg/mL Fosfomycin and 200 µg/mL Fosfomycin supplemented with 6 mM NAM. Each experimental category had at least three technical replicates. Experiment was done at least three times.

### Enzyme Kinetic by ATP Couple Assay

The kinetic studies for AmgK were performed with an ATP-coupled assay according to previously published protocol (62). Each reaction was made to a final volume of 100 µL, which contained 50 mM potassium phosphate buffer (pH = 7.5), 2 mM MgCl_2_, 1.0 mM PEP, 0.2 mM NADH, 4 U PK, and 3 U of LDH. In this assay, ATP was saturated at 2.5 mM, and the concentration of sugar substrates ranged from 0.01 mM to 2.5 mM. The reaction began with the addition of 1 ug of AmgK at 25 ^°^C and recorded for 3 minutes. The absorbance was measured every 5 seconds at 340 nm using a spectrophotometer (Eppendorf 6136). The linear region of the absorbance curve was determined to perform kinetic calculations. Collected data were processed and displayed to the Michaelis-Menten equation using GraphPad Prism 7.

### Time dependent MH-S invasion study with AzNAM labeled ***Legionella***

The *Legionella* invasion protocol was adapted from previous publications (31, 49). Briefly, a 24-well plate with sterile round cover glasses was treated with 500 µL of 0.1 mg/ mL poly-L-ornithine at 4 °C overnight. MH-S cells were washed twice with 500 µL of sterile PBS, seeded onto poly-L-ornithine coated round coverslips at a density of 100,000 cells per well in RPMI growth medium, and incubated overnight at 37 °C with 5% CO₂. The next day, MH-S cells were washed with RPMI medium without antibiotics and resuspended on 500 µL RPMI medium without antibiotics. *Legionella* cells were remodeled with the AzNAM probe as described earlier. After three hours of incubation with AzNAM, *Legionella* cells were washed twice with sterile PBS, resuspended to an OD₆₀₀ of ∼2.0, and 20 µL was added to each well containing MH-S cells, followed by centrifugation at 500 × g for 2 minutes and incubation at 37 °C with 5% CO₂ for 30, 60, 90, and 120 minutes. After incubation, the medium was replaced with 50 µg/ mL gentamicin in RPMI and incubated at 37 °C and 5% CO_2_ for 30 minutes to remove extracellular *Legionella*.

Cells were washed with 500 µL sterile PBS and fixed with 4% paraformaldehyde in PBS for 10 min at room temperature. Cells were washed three times with 400 µL PBS for five minutes on a rocker. To give access to intercellular *Legionella* of click reagents for reacting with Az handle, MH-S cells were permeabilized with 400 µL freshly prepared 1% Triton-X in PBS for 10 min. Then the cells were washed three times with 400 µL washing medium (0.2% tween-20 and 1.5% BSA in PBS) and resuspended on 400 µL of 0.1% BSA and 0.1% Triton-X solution in PBS for click reaction. 2 µL of 50mM CuSO_4_, 2 µL of 7 mM BTTAA, 2 µL of freshly prepared 12mg/ mL NaSc and 0.8 µL of 1 mg/mL Alk-488 were added sequentially and incubated at RT for 30 min in dark while shaking. Cells were stained with Phalloidin-TRITC (1:200) in 1.5% BSA in PBS for one hour after washing three times with the washing medium. After one hour of incubation, cells were washed three times and mounted on a glass slide with 4,6-diamidino-2-phenylindole (DAPI) (Invitrogen) and incubated overnight at RT in the dark. Next day, glass slides were sealed with clear nail polish to use for imaging with a confocal microscope (LSM800).

### Intracellular replication assay

Intracellular replication assay was adapted from previously published protocol (46, 71). In short, 10^5^ MH-S cells were seeded in 24 well plates maintaining 500 µL volume of RPMI medium and incubated overnight at 37 °C. Each well was washed three times with pre-warmed RPMI medium and resuspended in 450 µL RMPI. An overnight grown *Legionella*-pJB908 and *Legionella ΔamgK* cells were diluted to 2 × 10^6^ cells / mL in RPMI medium. From the diluted cell suspension, 50 µL of bacterial cells were added to get 10^5^ bacterial cells to maintain multiplicity of infection (MOI) 1. To enhance contact of bacterial cells to MH-S cells, the 24 well plates were centrifuged at 500g for 3 minutes. Plates were incubated for 2 hours, 24 hours, 48 hours and 72 hours at 37°C with 5% CO_2_. After the incubation, each well was washed 3 times with RPMI medium and then replaced with 500 µL of 0.05% digitonin in sterile PBS. MH-S cells were lysed with pipetting 20 times in 10 minutes. 100 µL of lysate was taken to make serial dilution in sterile PBS. The two hours of incubation period marks the infection time point, during which bacterial cells are expected to invade MH-S cells without sufficient time for replication. Therefore, this time point gave infectivity information while 24 hours to 72 hours incubation provided intracellular replication information. For 2 hours incubation cells were diluted to 10^0^ and 10^1^, for 24 hours incubation cells were diluted to 10^1^-10^2^, for 48 - 72 hours incubation cells were diluted to 10^3^ and 10^4^ . Cells were then spread on CYE-agar medium and incubated at 37 °C for four days to get colony count.

## Acknowledgments

We are thankful for support from the Delaware COBRE program, supported by a grant from the National Institute of General Medical Sciences (NIGMS 1 P30 GM110758 and 1 P20 GM104316-01A1). This work was supported by the NIH (U01 CA221230, R21 AI163949, R01 R01GM138599 (to CLG) and R01 GM132460 (to JMF)). We would like to thank Jeff Caplan and the Delaware Biotechnology Institute Bioimaging Center, where microscopy access was supported by grants from the NIH-NIGMS (P20 GM103446), the NIGMS (P20 GM139760) and the State of Delaware. Andor Dragonfly acquired with NIH-NIGMS (S10 OD030321). We would also like to thank William Trout, Shi Bai and the UD NMR facility, as well as PapaNii Asare-Okai, from the UD Mass Spectrometry Core Facility. C.L.G. thanks the Dreyfus foundation for support. S.N.H. would like to thank the NIH for support through the Chemistry-Biology Interface (CBI) training grant, T32GM133395. Figures were created using BioRender.

## Supporting information

**S1 Data: AmgK and MurU sequences of *P. putida* and *L. pneumophila***

**S1 Table. List of Strain and Plasmid Used**

**S1 Fig. Bioinformatic analysis of *L. pneumophila* proteins sequences.** (A) Protein homology of AmgK and MurU of *P. putida* with *L. pneumophila*. (B) AmgK and MurU of *P. putida* proteins distance tree with homolog of *L. pneumophila*.

**S3 Fig. Construction of *Legionella ΔamgK* using NEBuilder HiFI assembly.** (A) Schematic of the Allelic Exchange Substrate (AES) construction. (B) Five fragments were PCR-amplified for AES construction. Apapter_up, *lpg0295, thyA, lpg0297 and* Adapter_Down was 2054 bp, 676 bp, 972 bp, 1154 bp and 1029 bp respectively. (C) Shows PCR amplified HiFi assembled product from five fragments at 5876 bp. (D) Colony of *Legionella* Δ*amgK* grown after transformation with AES on the CYE-agar plate without thymidine. (E) Verification of *amgK* knockout with colony PCR with 1195 bp band indicative as positive colonies that replace *amgK* gene with *thyA* gene on the *Legionella* genome. *Legionella* genomic DNA was used as template for negative control and AES was used as template for positive control for colony PCR verification.

**S2 Table. List of primers used for *amgK* knockout**

**S3 Table. HiFi cloning conditions for *amgK* gene knockout**

**S4 Fig. Construction of *amgK* complement on *Legionella* Δ*amgK* in pMMB207c vector.** (A) Design of pMMB207c with *amgK* gene under *lac* operator. (B) Target gene *amgK* PCR amplification from genomic DNA of *Legionella*. (C) Colony PCR to verify the presence of *amgK* gene on HiFi Assembled pMMB207c vector in DH5α cells transformed with the plasmid on LB-Cm plate. (D) *Legionella ΔamgK-*pMMBc207c*-amgK* on CYE-agar plate with chloramphenicol. (E) Colony PCR for *Legionella ΔamgK-*pMMBc207c*-amgK* to verify the presence of *amgK* on *Legionella* Δ*amgK*.

**S4 Table. List of primers used for pMMB207C-*amgK* complement S5 Table. HiFi cloning conditions for *amgK* complement**

**S5 Fig. Structural alignment of AmgK and MurU of *Legionella* and *P. aeruginosa*.** Both AmgK PDB files were from predicted AlphaFold structure. While MurU of *Legionella* PDB file was from AlphaFold predicted structure, *P. aeruginosa* MurU (PDB ID: 8HHD) structure was available from PDB file from crystal structure study.

**S6 Fig. Construction of expression vectors for *E. coli* model system.** (A) pBBR-MCS-1-PA-KU HI vector with IPTG inducible *lac* promoter with SpcR gene. (B) Western blot expression test with 6xHis tag for 39 kDa and 25 kDa PA AmgK and MurU respectively. (C) Shows pBBR-MCS-1-Lp-KU-FLAG vector with IPTG inducible *lac* promoter with CamR gene. (D) Western blot expression test with FLAG tag for 42 kDa and 25 kDa *Legionella* AmgK and MurU respectively. **S6 Table. List of primers for E. coli *ΔmurQ* model system**

**S7 Table. HiFi Cloning Conditions for Expression vector construction for *E. coli* model system**

**S7 Fig. AmgK expression vector construction and purification.** (A) AmgK expression vector pGEX-AmgK. (B) Coomassie stained SDS-PAGE gel image of purified AmgK (∼38 kDa).

**S8 Table. List of primers used for AmgK protein expression on pGEX vector S9 Table. HiFi cloning conditions for expression vector carrying *amgK***

**S8 Fig. Time-point study of MH-S invasion with AzNAM Labeled *Legionella*.** The figure shows images of MH-S cells co-cultured with AzNAM labeled *Legionella*. Az-NAM labeled *Legionella* cells were co-culture with MH-S cells for 30 min, 60 min, 90 min and 120 min and fixed prior to the CuAAC click labeling (*Legionella*, Alk-488, Green) and subsequent labeling with Phalloidin-TRITC (F-actin, Red) and DAPI (nucleus, blue). Images were taken on a Zeiss LSM800 confocal microscope with Z-stack of multiple planes and combined with orthogonal projection with scale bars =10 μm. Images are representative of a minimum of three fields viewed per replicate with at least three technical replicates, and experiments were conducted in at least three biological replicates.

**S9 Fig. Time-course of MH-S invasion with NAM labeled *Legionella*.** The figure shows images of MH-S cells co-cultured with NAM labeled *Legionella* as negative control for labeling. NAM labeled *Legionella* cells were co-culture with MH-S cells for 30 min, 60 min, 90 min and 120 min and fixed prior to the CuAAC click labeling (*Legionella*, Alk-488, Green) and subsequent labeling with Phalloidin-TRITC (F-actin, Red) and DAPI (nucleus, blue). Images were taken on a Zeiss LSM800 confocal microscope with Z-stack of multiple planes and combined with orthogonal projection with scale bars =10 μm. Images are representative of a minimum of three fields viewed per replicate with at least three technical replicates, and experiments were conducted in at least three biological replicates.

**S10 Fig. Mass spec analysis of reduced AzNAM incorporation in the presence of FerricNitrate in the AYE medium.** Mass spectrometry analysis to verify NH_2_NAM incorporation on lysozyme digested disaccharide fragments following remodeling with AzNAM. In the bottom it shows the simulation of three isotopes with mass and the top panel shows the relative abundance of each isotope in the sample.

