## Supplementary material for "The *Legionella pneumophila* peptidoglycan recycling kinase, AmgK, is essential for survival and replication inside host alveolar macrophages": SI Figures and Data

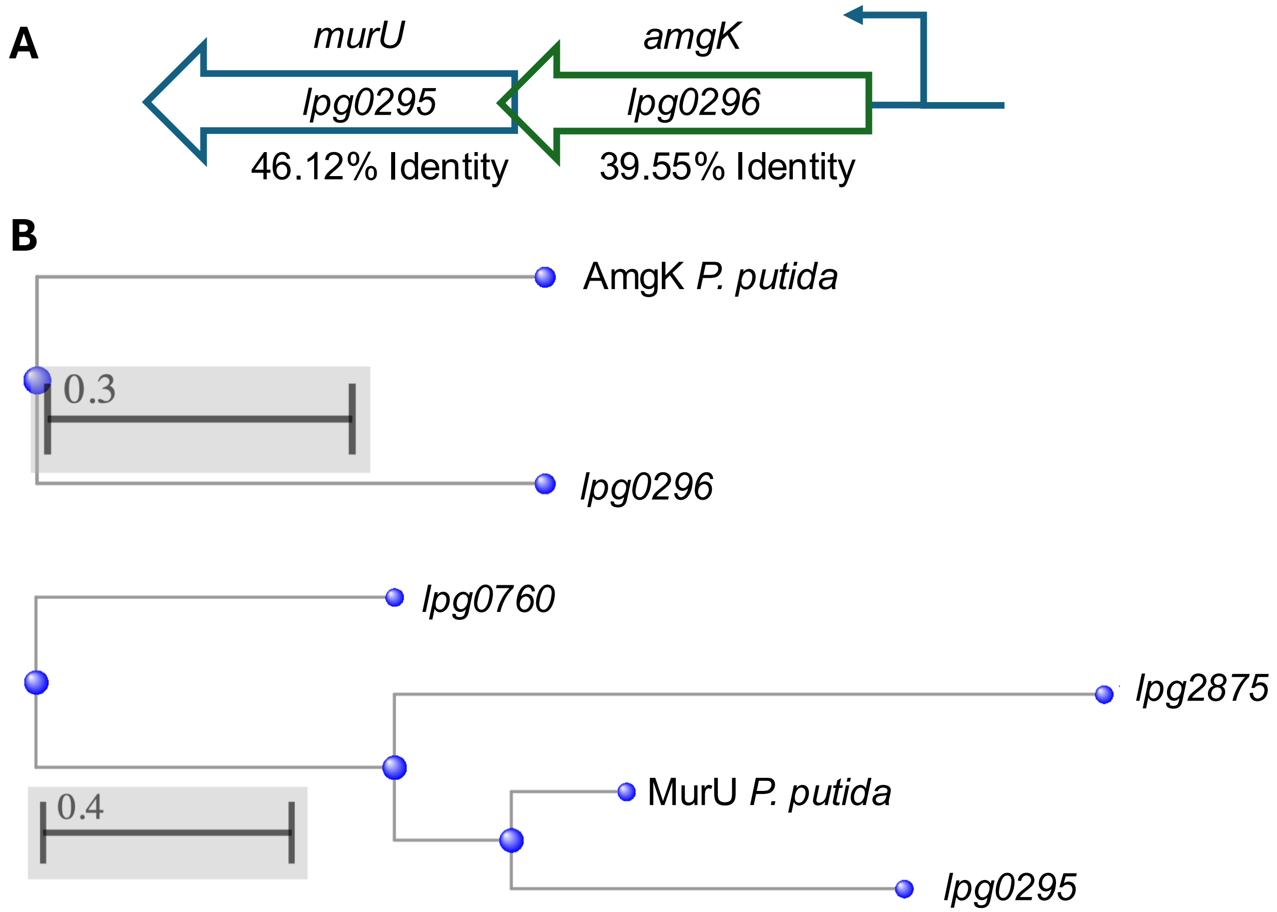
A
murU
lpg0295
46.12% Identity
amgK
lpg0296
39.55% Identity
B
AmgK P. putida
0.3
lpg0296
lpg0760
lpg2875
MurU P. putida
0.4
lpg0295

**S1 Fig**

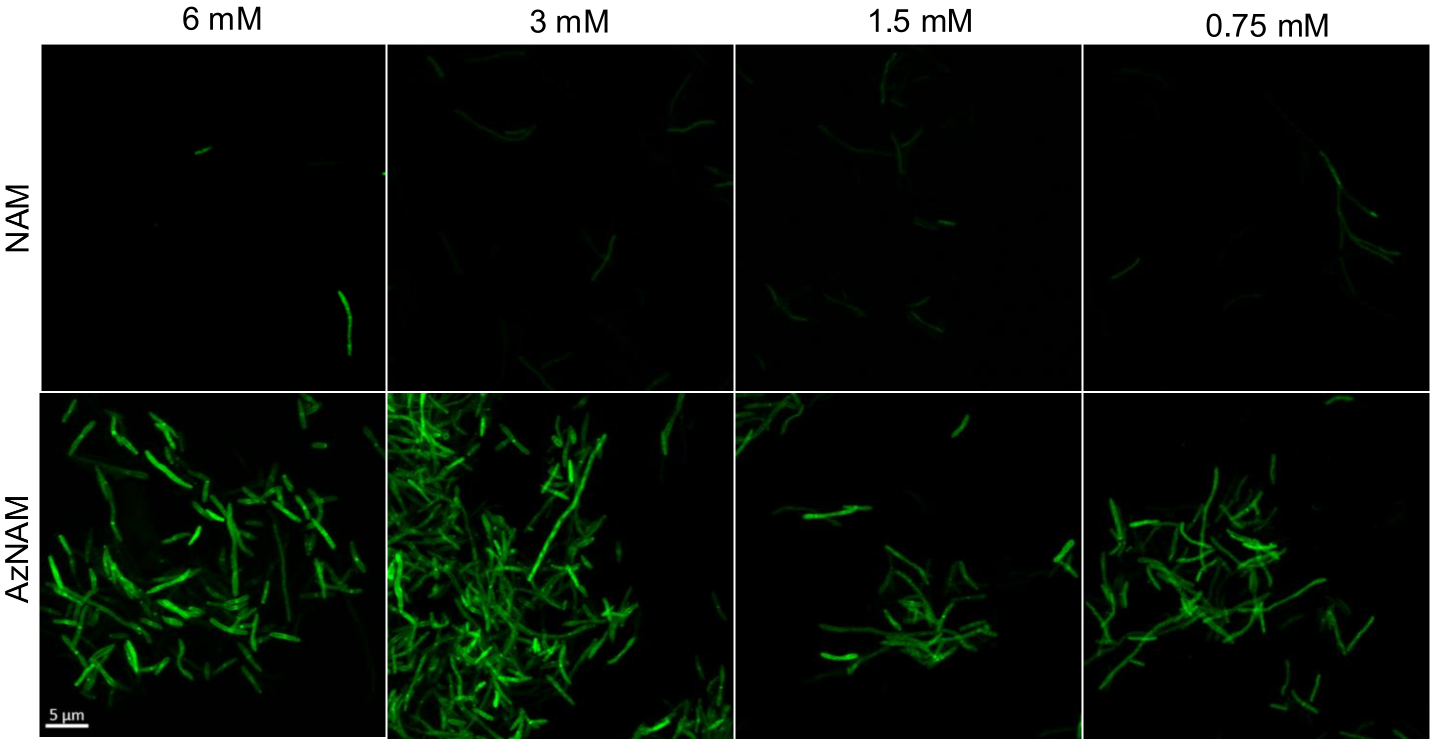


**S2 Fig**

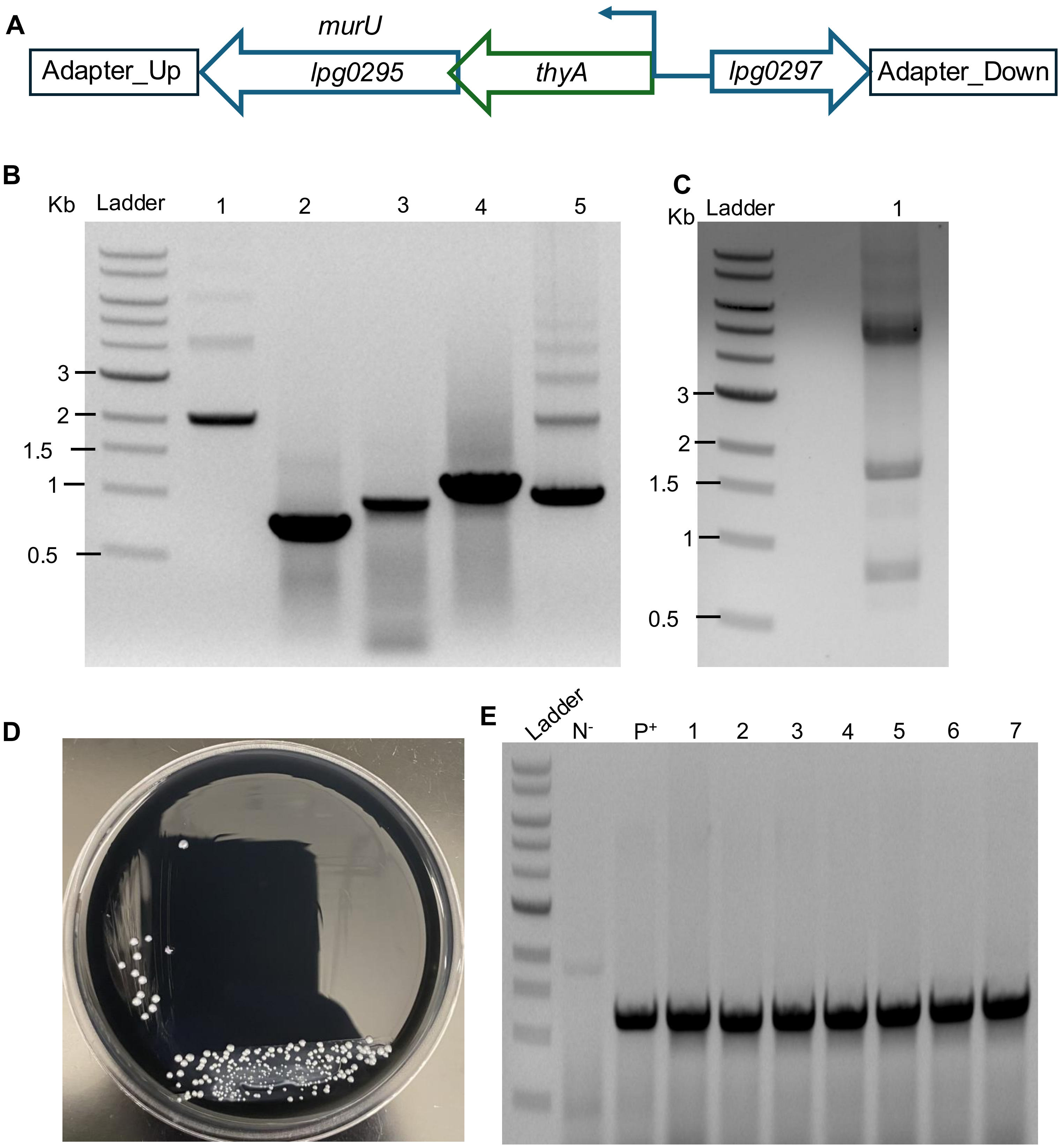
A
murU
Adapter_Up
lpg0295
thyA
lpg0297
Adapter_Down
B
Kb
Ladder
1
2
3
4
5
3
2
1.5
1
0.5
C
Kb
Ladder
1
3
2
1.5
1
0.5
D
E
Ladder
N-
P+
1
2
3
4
5
6
7

**S3 Fig**

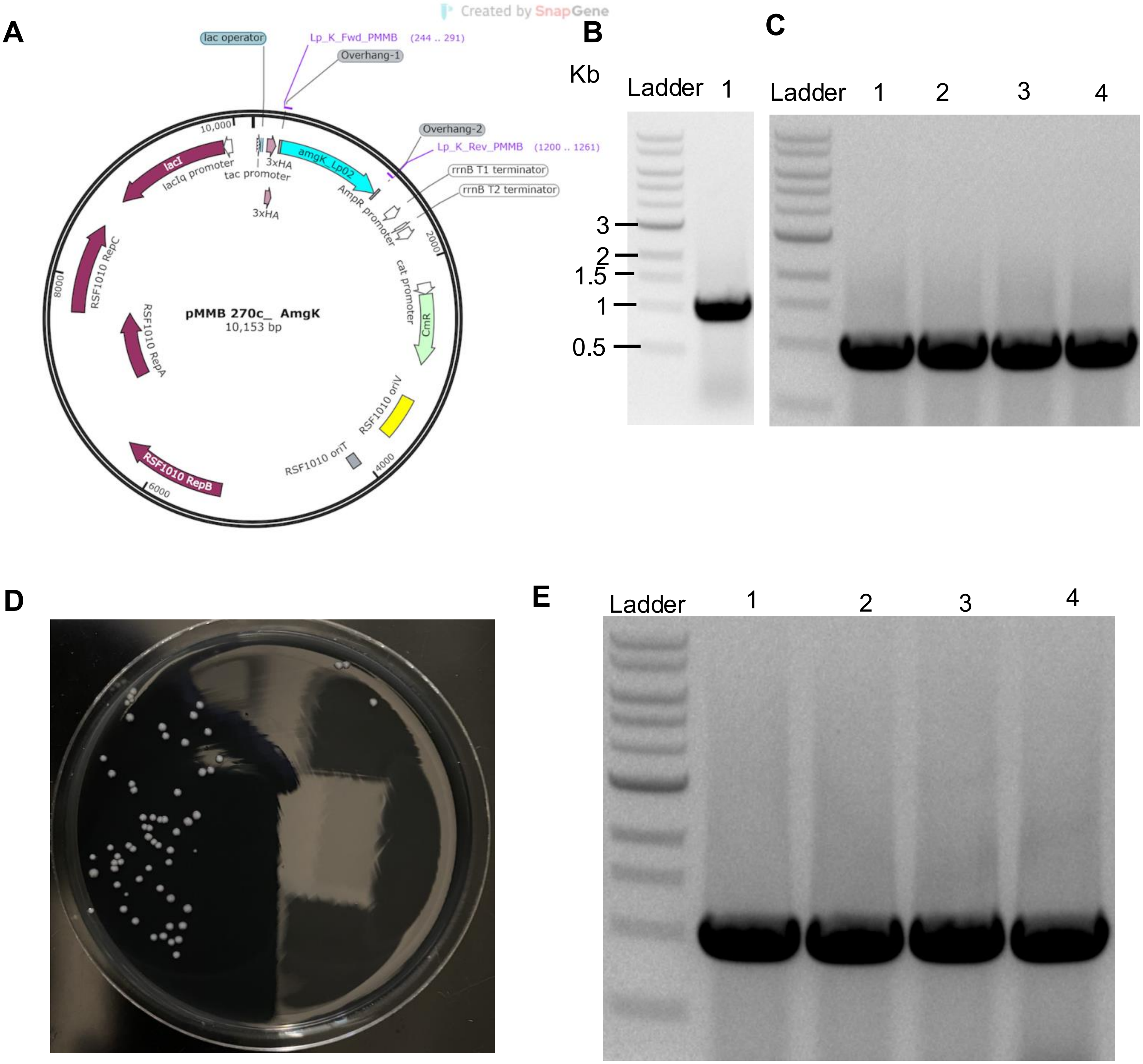

A
Created by SnapGene
lac operator
Lp_K_Fwd_PMMB (244 .. 291)
Overhang-1
Overhang-2
Lp_K_Rev_PMMB (1200 .. 1261)
rrnB T1 terminator
rrnB T2 terminator
10,000
lacI
lacIq promoter
tac promoter
3xHA
amgK_Lp02
AmpR promoter
2000
cat promoter
CmR
RSF1010 RepC
8000
RSF1010 RepA
pMMB 270c_ AmgK
10,153 bp
RSF1010 oriV
RSF1010 oriT
4000
RSF1010 RepB
6000
B
Kb
Ladder
1
3
2
1.5
1
0.5
C
Ladder
1
2
3
4
D
E
Ladder
1
2
3
4


**S4 Fig**

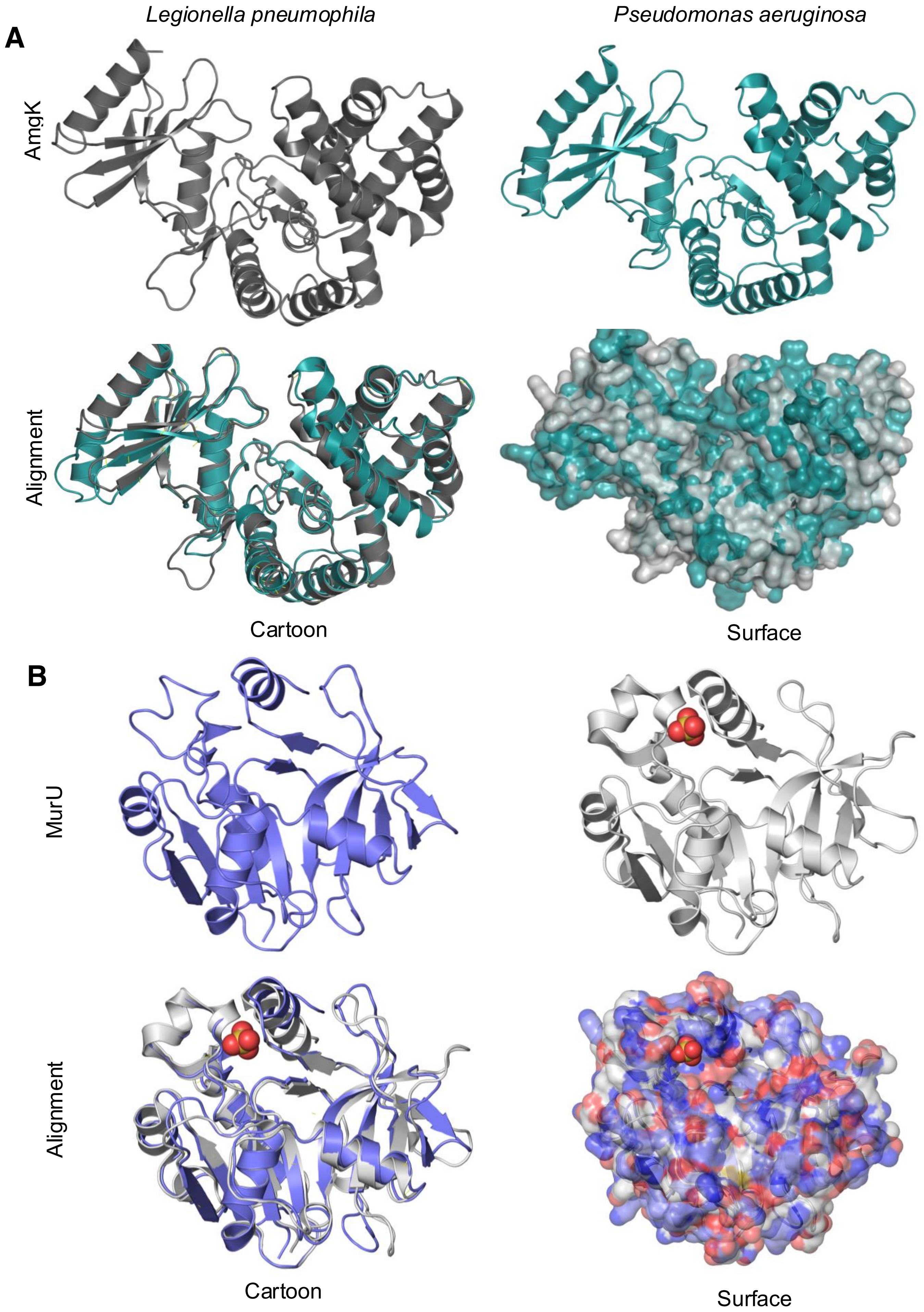
Legionella pneumophila
Pseudomonas aeruginosa
A
AmgK
Alignment
Cartoon
Surface
B
MurU
Alignment
Cartoon
Surface

**S5 Fig**

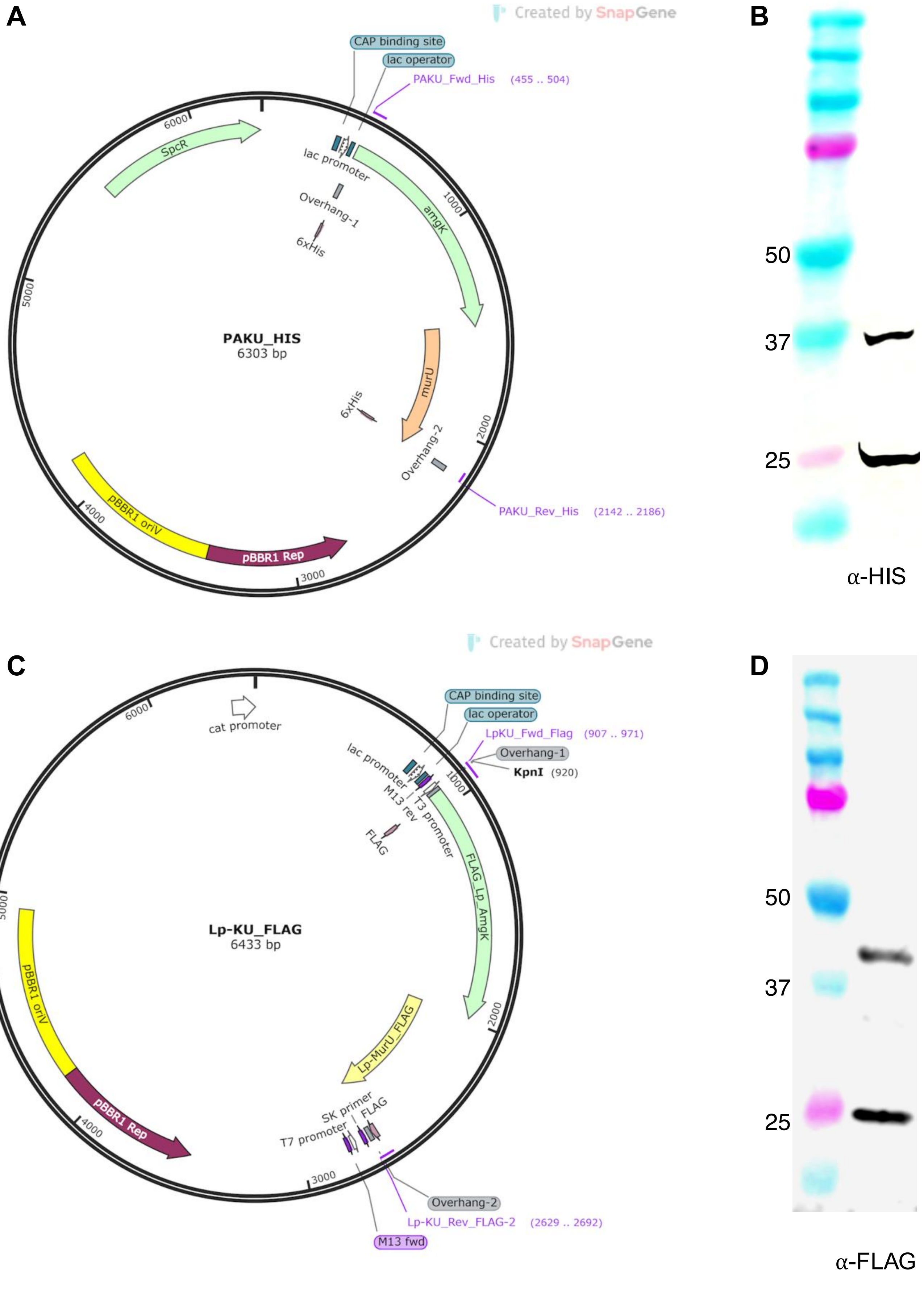
A
Created by SnapGene
CAP binding site
lac operator
PAKU_Fwd_His (455 .. 504)
SpcR
lac promoter
Overhang-1
6xHis
amgK
1000
6000
5000
PAKU_HIS
6303 bp
murU
6xHis
Overhang-2
2000
PAKU_Rev_His (2142 .. 2186)
pBBR1 oriV
4000
pBBR1 Rep
3000
B
50
37
25
α-HIS
C
Created by SnapGene
CAP binding site
lac operator
LpKU_Fwd_Flag (907 .. 971)
Overhang-1
KpnI (920)
cat promoter
6000
lac promoter
M13 rev
T3 promoter
FLAG
1000
FLAG_Lp_AmgK
5000
Lp-KU_FLAG
6433 bp
pBBR1 oriV
Lp-MurU_FLAG
2000
pBBR1 Rep
4000
SK primer
FLAG
T7 promoter
3000
Overhang-2
Lp-KU_Rev_FLAG-2 (2629 .. 2692)
M13 fwd
D
50
37
25
α-FLAG

**S6 Fig**

**A**

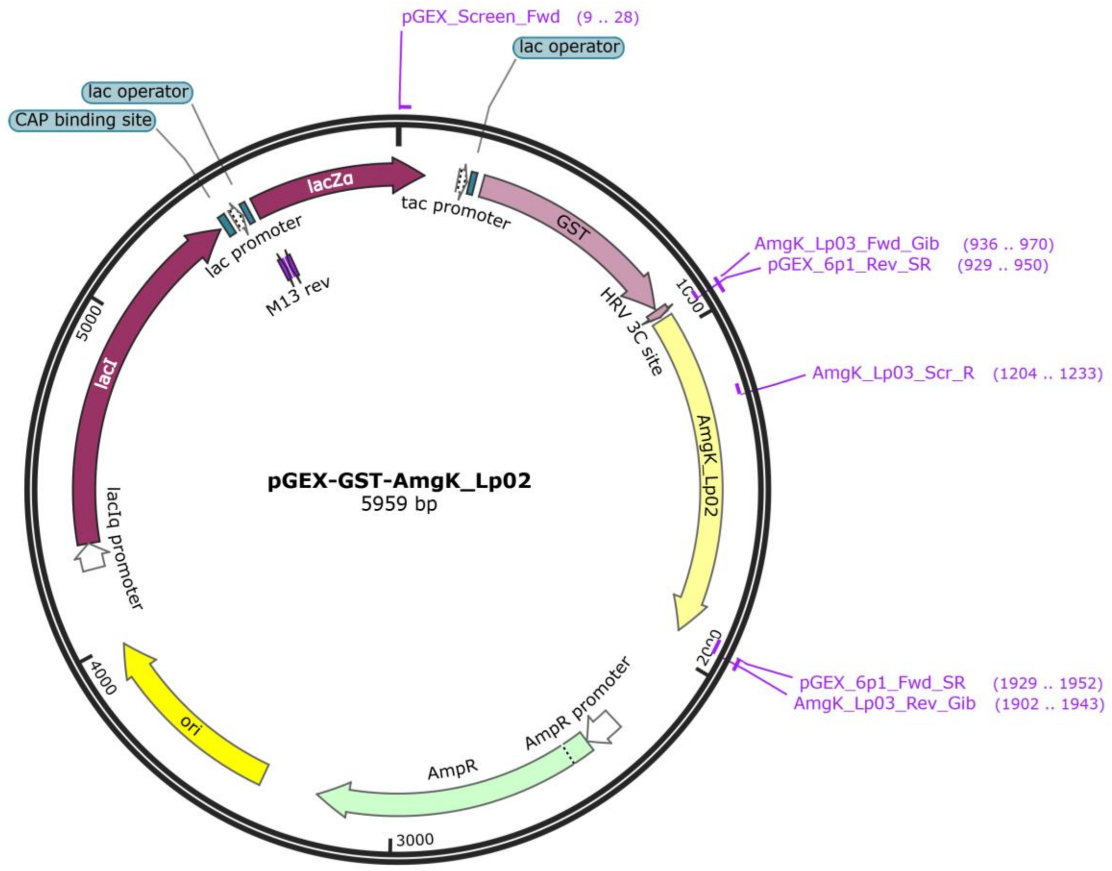
pGEX_Screen_Fwd (9 .. 28)
lac operator
lac operator
CAP binding site
lacZα
tac promoter
GST
lac promoter
M13 rev
AmgK_Lp03_Fwd_Gib (936 .. 970)
pGEX_6p1_Rev_SR (929 .. 950)
HRV 3C site
AmgK_Lp03_Scr_R (1204 .. 1233)
5000
lacI
pGEX-GST-AmgK_Lp02
5959 bp
AmgK_Lp02
lacIq promoter
pGEX_6p1_Fwd_SR (1929 .. 1952)
AmgK_Lp03_Rev_Gib (1902 .. 1943)
4000
AmpR promoter
ori
AmpR
3000

**B**

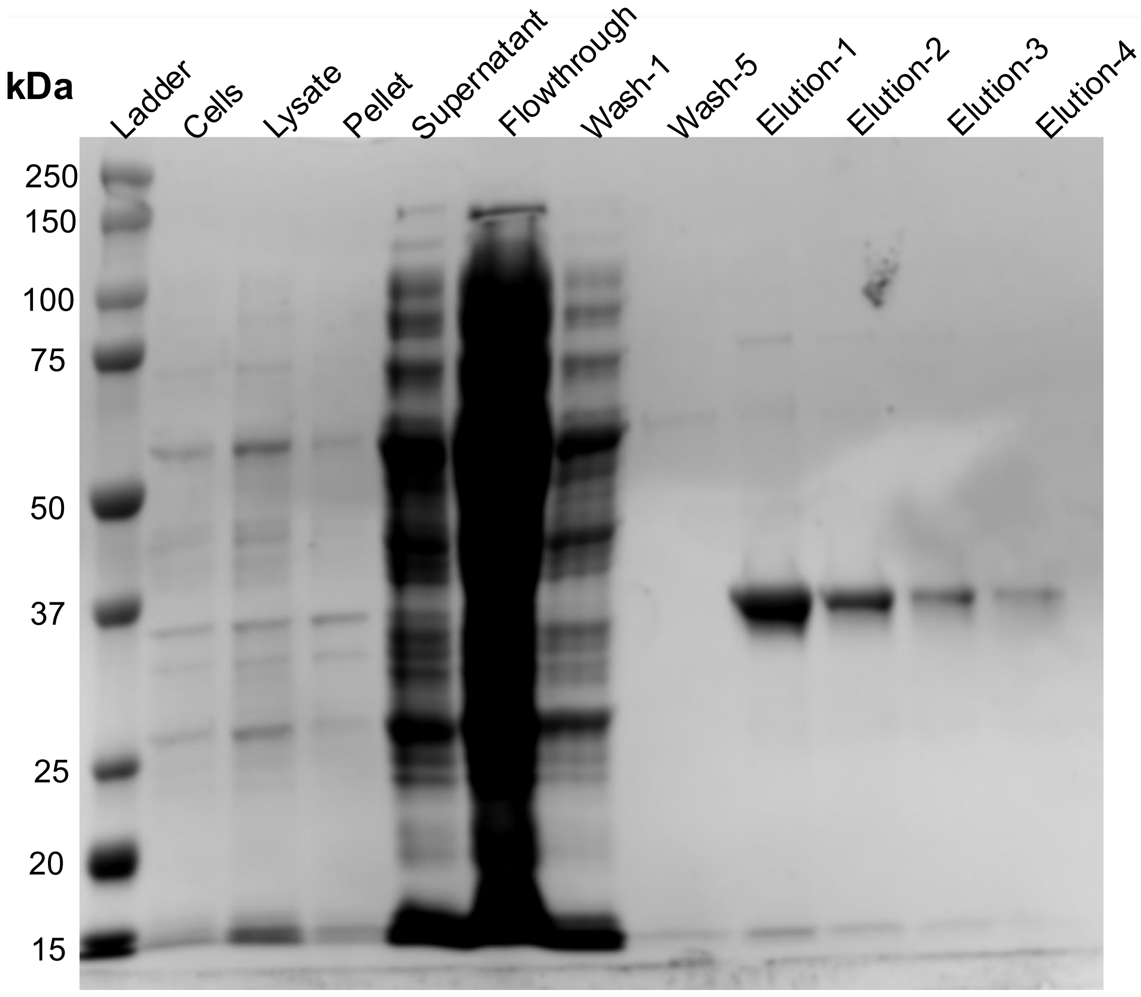
kDa
Ladder
Cells
Lysate
Pellet
Supernatant
Flowthrough
Wash-1
Wash-5
Elution-1
Elution-2
Elution-3
Elution-4
250
150
100
75
50
37
25
20
15

**S7 Fig**

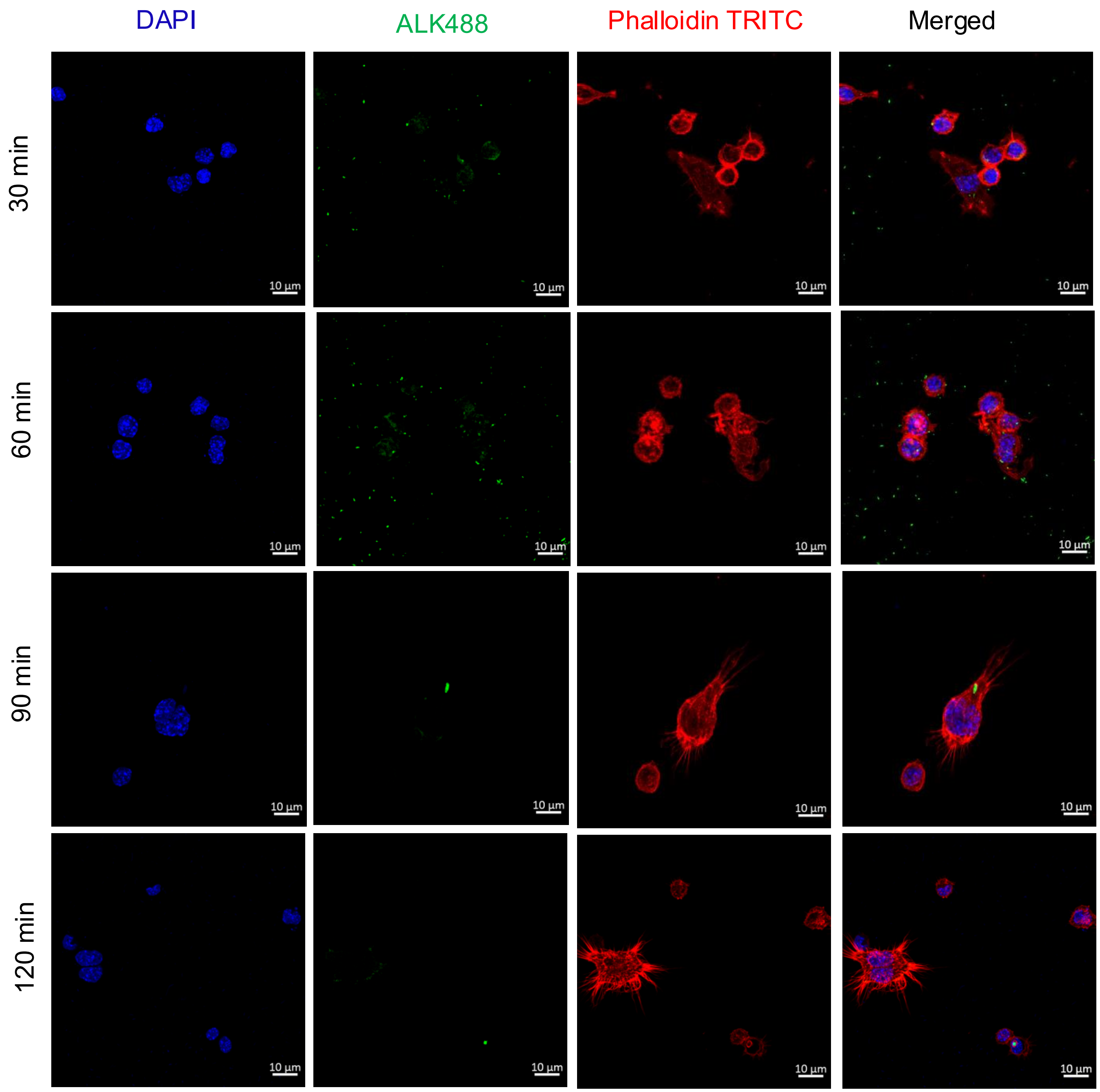
DAPI
ALK488
Phalloidin TRITC
Merged
30 min
60 min
90 min
120 min
10 µm

**S8 Fig**

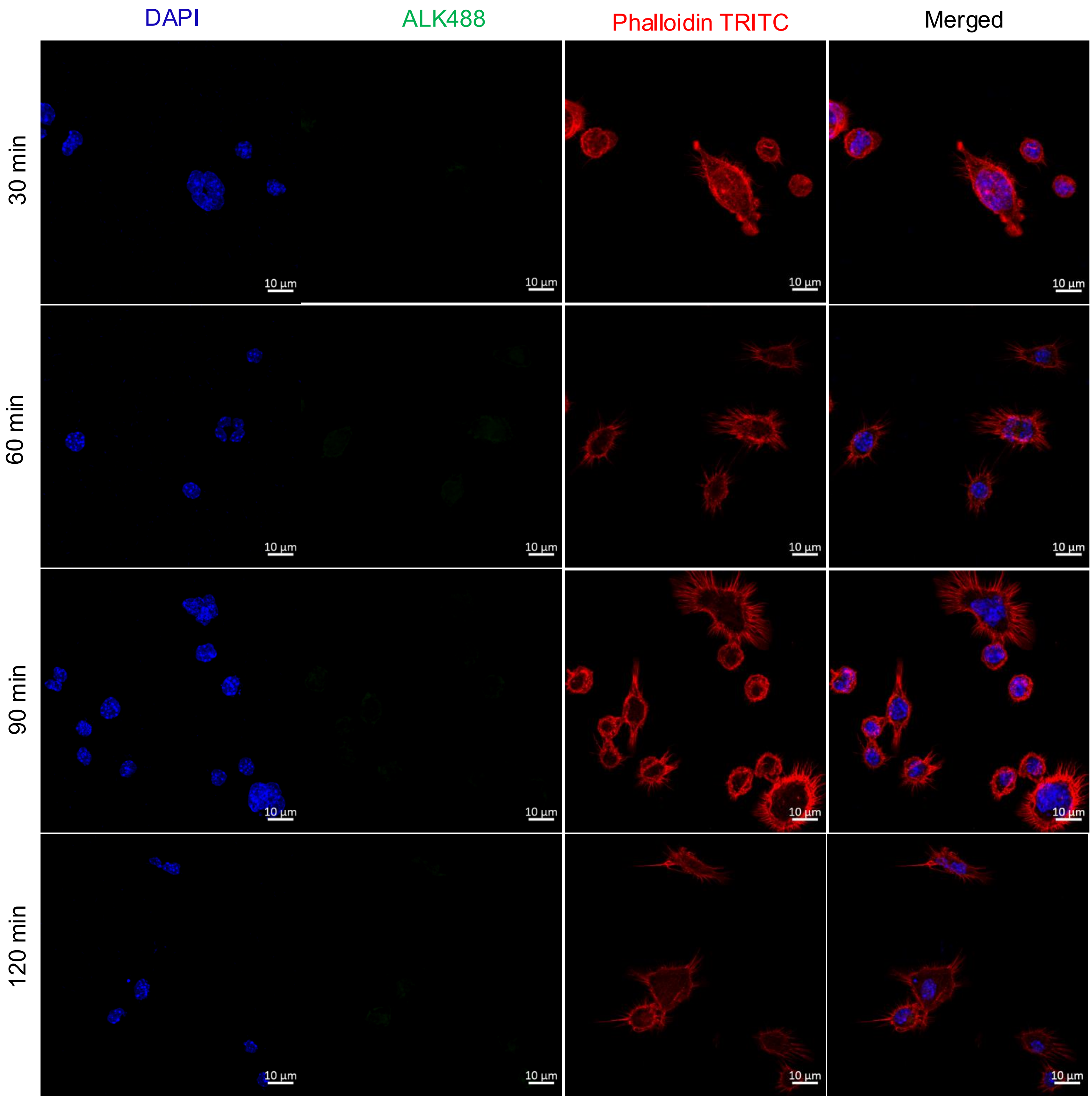
DAPI
ALK488
Phalloidin TRITC
Merged
30 min
60 min
90 min
120 min
10 µm

**S9 Fig**

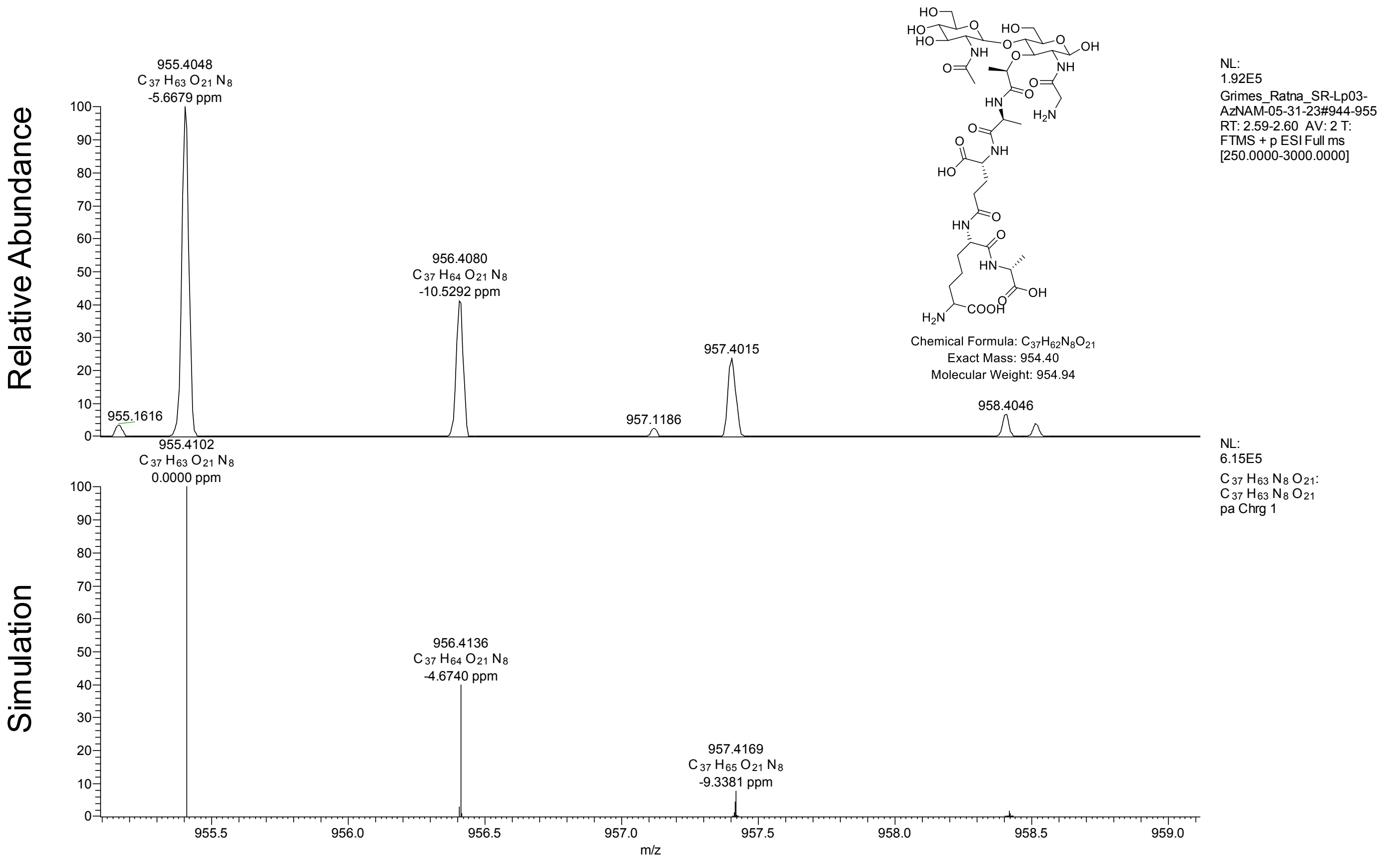

Relative Abundance
Simulation
955.4048
C 37 H63 O21 N8
-5.6679 ppm
955.1616
956.4080
C 37 H64 O21 N8
-10.5292 ppm
957.1186
957.4015
958.4046
955.4102
C 37 H63 O21 N8
0.0000 ppm
956.4136
C 37 H64 O21 N8
-4.6740 ppm
957.4169
C 37 H65 O21 N8
-9.3381 ppm
m/z
Chemical Formula: C37H62N8O21
Exact Mass: 954.40
Molecular Weight: 954.94
NL:
1.92E5
Grimes_Ratna_SR-Lp03-AzNAM-05-31-23#944-955
RT: 2.59-2.60 AV: 2 T:
FTMS + p ESI Full ms
[250.0000-3000.0000]
NL:
6.15E5
C 37 H63 N8 O21:
C 37 H63 N8 O21
pa Chrg 1


**S10 Fig**

Protein sequence of AmgK and MurU of *P. putida* were used as quarry sequences to BLAST with whole protein sequence of Lp02 as subject.

### *amgK* (*lpg0296*) sequence from Lp_Philadelphia 1.

atgcatgaaagagaaaacgcactgaaagagtggttagctcaaacgattcaacaaaaagactttgtcctccttcctttagcaggagatg
cgagttttagaaggtattttcgcattcaatataatggattaactcaagtagttatggacgccccacctgaaaaggaaaatattgaacctttc
ctccatatcgctaacgttcttgataaaattaaaataccagtccccgatattctggcaataaataaaagagaaggtttcctgttactaagcg
atttaggggatcaattattattaaataagctgaatcatgaaacagtggataattattataaccaggcaatgaatttgctattacaaattcaa
aaatgtcccactgacgatcctgaacttcccttatttgacaaaagtttcatgcttaaagaaatgaatttatgtttggaatggtttttgaaagctta
cctttctcttgatttaaaacaagacgaattacaactttttcagcataccatcgagtggatcgcaagtgaagtagcgactcaaccaagagtt
ttcattcacagggactaccattcaagaaatctgatgctagtcaaaaataaggacagctgtttagcaactattgattttcaggatgctatgct
aggtcctgttacctatgatttggtttctttactcaaagattgttatatttcatggccaagagaaaaagttttagaatgggtaacctattttcatga
acaatccccccctcgccagtatttactctctaaccgattttatccgggcctttgatctatgtgggttacaaaggcatttaaaagtgttaggtgttt
tttgccgtttatatttgcgtgataataaagcaggatatcttggggacttgcctctcacgttaaaatatgcattagaatgcgcagaaacatatg
aagaattacacccatttttttaattttctgcaaaaacgagtttatttaccatga

### AmgK (WP_015444878.1) protein sequence from Lp_Philadelphia 1.

MHERENALKEWLAQTIQQKDFVLLPLAGDASFRRYFRIQYNGLTQVVMDAPPEKENIEPFLHIA
NVLDKIKIPVPDILAINKREGFLLLSDLGDQLLLNKLNHETVDNYYNQAMNLLLQIQKCPTDDPEL
PLFDKSFMLKEMNLCLEWFLKAYLSLDLKQDELQLFQHTIEWIASEVATQPRVFIHRDYHSRNL
MLVKNKDSCLATIDFQDAMLGPVTYDLVSLLKDCYISWPREKVLEWVTYFHEQSPLASIYSLTD
FIRAFDLCGLQRHLKVLGVFCRLYLRDNKAGYLGDLPLTLKYALECAETYEELHPFFNFLQKRV
YLP

### *murU* (*lpg0295*) gene sequence from Lp_Philadelphia 1.

atgaaaaccgctatgattcttgcagcaggacgtggagaacgattacgccctttaaccgacaaaatgcctaaagcactttgtacggtaa
gaaacaaacccttaattgaacaccatatcattaacctcgccaatgccggctttgaacgattgataatcaatcatgcctacctgggcgga
caaattcgccaatatataggaaatggaaaacaatggggcttaaatgttatctattctccagaaccgccaggaggccttgaaacaggg
ggagggattgtgaacgccttgcctttgctgggcgaagagccatttttaacagtaaatgccgacatttatactgattttgattttgccaagctc
caactcaaaaatattgatacatttcatctgcttttaattcccaaaaatccttctttgcttcatcatggcgattttggtttaatcaatggatcccact
taaccaatacaaatcaggcttacactttttcaggtattgcctgctataacccaaaggtttttgccaactgcagacaaggaaggtattctgtt
accccttttactcagaaaatatgtggaacaaaatagtgccactggaagtgttcactcaggaatttggtatgacataggctctttcgataga
ctgcaggcagcaaatcagttcacctaa

### MurU (WP_010946056.1) protein sequence from Lp_Philadelphia 1.

MKTAMILAAGRGERLRPLTDKMPKALCTVRNKPLIEHHIINLANAGFERLIINHAYLGGQIRQYIG
NGKQWGLNVIYSPEPPGGLETGGGIVNALPLLGEEPFLTVNADIYTDFDFAKLQLKNIDTFHLLLI
PKNPSLLHHGDFGLINGSHLTNTNQAYTFSGIACYNPKVFANCRQGRYSVTPLLRKYVEQNSA
TGSVHSGIWYDIGSFDRLQAANQFT

**S1 Table. List of Strain and Plasmid Used**

| Strains or Plasmids | Key Characters | References |
|---|---|---|
| *Legionella pneumophila* (Lp02) | Thymidine Auxotroph | (1) |
| *Legionella pneumophila ΔamgK* | Can Grow without Thymidine | This Study |
| *Legionella pneumophila ΔamgK*-*pamgK* | $Cam^R$ | This Study |
| DH5α | Plasmids Construction | (2) |
| BL21-pGEX-GST-amgK | Protein Expression | This Study |
| *E. coli ΔmurQ* | $Kan^R$ | |
| pDG1662 | $Amp^R$, $Spc^R$ | (3) |
| pMMB207c 4xHA | $Cam^R$ | Dr. Gunnar Schroeder Lab (Queen's University Belfast) |
| pMMB207c-amgK | $Cam^R$ | This Study |
| pJB908 | thyA Gene | (4) |
| pBBR-Lp-KU-FLAG | $Cam^R$ | This Study |
| pBBR-PA-KU-6His | $Spc^R$ | This Study |
| pGEX-6P1-amgK (GST) | $Amp^R$ | This Study |
| MH-S | Murine Alveolar Macrophage | ATCC |

**S1 Table. List of Strain and Plasmid Used**

| Strains or Plasmids | Key Characters |
|---|---|
| *Legionella pneumophila* (Lp02) | Thymidine Auxotroph |
| DH5α | Plasmids Construction |
| *E. coli ΔmurQ* | $Kan^R$ |
| pDG1662 | $Apm^R$, $Spc^R$ |
| pMMB207c | $Cam^R$ |
| MH-S | Murine Alveolar Macrophage |

**S3 Table. HiFi Cloning Conditions for *amgK* gene knockout**

| Constructs name | Gibson assembly calculation | Final size |
|---|---|---|
| amgK_KO_AES | pDG62 Up 101.6 ng (2054 bp) : amgK_Down 37.8 ng (767 bp) : thyA 47.90 ng (972 bp) : amgK Up 57.36 ng (1164 bp) : pDG1662 Down 50.87 bp (1029 bp) [ 1:1 ratio with 80 fmol DNA fragments ] | 5876 bp |

**S4 Table. List of primers used for pMMB207C-*amgK* complement**

| Primes Name | Sequence 5'-3' | TA Phusion | PCR size |
|---|---|---|---|
| Lp_K_Fwd_PMMB | gcgcgggtacccggggatccatgcatgaaagagaaaacg cactgaaag | 68 ° C | 1018 bp |
| Lp_K_Rev_PMMB | ccgccaaaacagccaagctttcatggtaaataaactcgttttt gcagaaaattaaaaaatgg | | |

**S5 Table. HiFi cloning conditions for *amgK* complement**

| Constructs Name | Gibson Assembly Calculation | Final size |
|---|---|---|
| pMMB207c_amgK | pMMB (17.70 fmol): Lp amgK (35.39 fmol) = 100 ng (9175 bp): 22.19 ng (1018 bp) [1:2 ratio of Vector vs insert] | 10153 bp |

**S6 Table. List of primers for *E. coli ΔmurQ* model system**

| Primes Name | Sequence 5'-3' | TA Q-5 pol | PCR size |
|---|---|---|---|
| MCS1_Fwd | aagcttgatatcgaattcctgcagccc | 72 ° C | 4682 bp |
| MCS-1_Rev_KpnI | tggtacccagcttttgttccctttagtgag | | |
| LpKU_Fwd_Flag | caaaagctgggtaccaATGGATTACAAGGATGACGACGATAAATTACTCTTTATTAAGATGGgag | 58 ° C | 1786 bp |
| Lp-KU_Rev_FLAG-2 | ggaattcgatatcaagcttTCACTTATCGTCGTCATCCTTGTAATCggtgaactgatttgctgc | | |
| pBR_Fwd_His | catcaccatcaccatcactaggcgttaatattttg | 71 ° C | 4621 bp |
| pBR_Rev_His | gtgatggtgatggtgatgcatagctgtttc | | |
| PAKU_Fwd_His | agctatgcatcaccatcaccatcacTCTGATGATGCCCGTTTCCAGCAGC | 71 ° C | 1732 bp |
| PAKU_Rev_His | acgcctagtgatggtgatggtgatgGGCGTGCTCCGCCAGCAATC | | |

**S7 Table. HiFi cloning conditions for expression vector construction for *E. coli* model system**

| Constructs Name | Gibson Assembly Calculation | Final size |
|---|---|---|
| EQLPKU_FLAG | pBRMCS1 : LpKU_FLAG = 100 ng (4682 bp) : 76 ng (1786 bp) [ 1:2 ratio of Vector vs insert] | 6433 bp |
| EQPAKU_HIS | pBRMCS1_SpcR : PAKU_HIS = 100 ng (4621 bp) : 75 ng (1786 bp) [ 1:2 ratio of Vector vs insert] | 6303 bp |

**S8 Table. List of primers used for AmgK protein expression on pGEX vector**

| Primes Name | Sequence 5'-3' | TA Phusion | PCR size |
|---|---|---|---|
| pGEX_6p1_Fwd_SR | GAATTCCCGGGTCGACTCGAGCGG | 72 ° C | 4981 bp |
| pGEX_6p1_Rev_SR | GGATCCCAGGGGCCCCTGGAAC | | |
| AmgK_Lp03_Fwd_Gib | GGGCCCCTGGGATCCatgcatgaaagagaaaacgc | 61 ° C | 1008 bp |
| AmgK_Lp03_Rev_Gib | TCGACCCGGGAATTCtcatggtaaataaactcgttttgcag | | |
| pGEX_Screen_Fwd | GACTGCACGGTGCACCAATG | 67 ° C | 1225 bp |
| AmgK_Lp03_Scr_R | gatcccctaaatcgcttagtaacaggaaac | | |

**S9 Table. HiFi cloning conditions for expression vector with *amgK***

| Constructs name | Gibson assembly calculation | Final size |
|---|---|---|
| pGEX_amgK | pGEX (32.59 fmol): amgK (65.19 fmol) = 100 ng (4981 bp) : 40.48 ng (1008 bp) [ 1:2 ratio of Vector vs insert] | 5959 bp |
